# The Glucose Transporter GLUT3 Controls Regulatory T Cell Function

**DOI:** 10.64898/2026.03.26.714439

**Authors:** Katrin Sinning, Miriam Eckstein, Xiufeng Zhao, Alexandra Freitag, Mathias Rosenfeldt, Sophia M. Hochrein, Martin Vaeth

**Affiliations:** Würzburg Institute of Systems Immunology, Max Planck Research Group, Julius-Maximilians-University of Würzburg, Würzburg, Germany; Institute of Pathology, Julius-Maximilians-University of Würzburg, Würzburg, Germany

## Abstract

Regulatory T (Treg) cells are central mediators of immune tolerance and are generally considered to rely predominantly on mitochondrial metabolism rather than glucose-driven glycolysis. To define the role of glucose metabolism in Treg cells, we investigated the contribution of the hexose transporters GLUT1 and GLUT3. Genetic ablation of GLUT1 in T cells or selectively in Treg cells had minimal impact on Treg cell numbers, phenotype or immune homeostasis, indicating that GLUT1 is largely dispensable in this lineage. By contrast, deletion of GLUT3 in T cells resulted in a marked reduction in Treg cell numbers. However, it remained unclear whether this reduction reflected diminished IL-2 production by GLUT3-deficient conventional T cells or a cell-intrinsic requirement for GLUT3 in Treg cells. To investigate this, we generated mice with Treg cell-specific deletion of GLUT3. These animals developed severe systemic inflammation accompanied by lethal cellular and humoral autoimmunity. Mechanistically, GLUT3-deficient Treg cells exhibited reduced glycolytic activity and mitochondrial respiration, leading to impaired suppressive function and defective effector and follicular Treg cell differentiation. Collectively, our findings demonstrate a non-redundant requirement for GLUT3 in Treg cell metabolic fitness and immune regulation, refining the prevailing view that Treg cells operate largely independently of glucose metabolism. Our data further suggest that therapeutic strategies targeting glucose uptake and glycolysis in autoimmune and inflammatory diseases should account for potential adverse effects on Treg cell-mediated immune tolerance.

## INTRODUCTION

T regulatory (Treg) cells are essential mediators of immune homeostasis and play a crucial role in preventing autoimmunity. They are defined by expression of the transcription factor forkhead box P3 (Foxp3), which acts as a master regulator of their development and suppressive function, and by high levels of the interleukin-2 receptor α chain (CD25), enabling efficient interleukin-2 (IL-2) signaling required for their survival and functional stability (1). Through constitutive CD25 expression, Treg cells effectively compete for IL-2, a mechanism that both sustains Treg populations and limits the expansion of conventional effector T cells (2, 3). Loss of Foxp3 expression results in impaired Treg differentiation and function, leading to a breakdown of immune tolerance (1, 4). In mice, this is exemplified by the fatal autoimmune syndrome observed in “Scurfy” mice, which harbor a spontaneous loss-of-function (LoF) mutation in the *Foxp3* gene and exhibit widespread multiorgan inflammation and severe tissue pathology (5–7). This phenotype closely mirrors the human IPEX syndrome (immune dysregulation polyendocrinopathy enteropathy X-linked syndrome), a condition caused by LoF mutations in *FOXP3* that generally requires treatment by hematopoietic stem cell transplantation (8, 9). Together, these observations underscore the indispensable role of Foxp3⁺ Treg cells in maintaining immune tolerance and preventing systemic autoimmunity.

Although Foxp3 is indispensable for Treg cell lineage specification and suppressive function, it is not sufficient to fully establish the transcriptional, epigenetic and functional program of thymus-derived Treg cells. Ectopic expression of Foxp3 in conventional CD4⁺ T cells is insufficient to fully recapitulate the Treg-specific transcriptional and epigenetic landscape (4, 10), highlighting the requirement for additional regulatory mechanisms to establish and maintain the identity of Foxp3⁺ Treg cells.

Among the mechanisms governing T cell fate and effector function, cellular metabolism has emerged as a critical determinant of lymphocyte activation and differentiation (11–13). Unlike conventional effector T cells, which predominantly rely on aerobic glycolysis upon antigenic stimulation to support proliferation and effector functions, Foxp3⁺ Treg cells exhibit a distinct metabolic program, characterized by mitochondrial respiration and lipid oxidation (14–19). Consequently, multiple studies have demonstrated that mitochondrial biogenesis and oxidative phosphorylation (OxPhos) are essential for Treg cell homeostasis and function (20, 21), cementing the view that Treg cells rely far less – or not at all – on extracellular glucose and aerobic glycolysis. These observations are further corroborated by evidence that Foxp3 acts as a “metabolic gatekeeper”, restraining the expression of glucose transporters and glycolytic genes (18, 22). Conversely, inducible Foxp3 deletion in Treg cells unleashes glycolytic activity, accompanied by impaired suppressive function (23), suggesting that excessive glycolysis compromises both the lineage fidelity and functional competence of Treg cells.

Challenging the view that Foxp3^+^ Treg cells are exclusively reliant on mitochondrial respiration, recent studies have demonstrated that glucose metabolism plays a significant role in supporting Treg homeostasis, migratory capacity and effector differentiation, especially in the context of inflammation (22, 24–29). The first step in the initiation of aerobic glycolysis involves the uptake of extracellular glucose. This process is facilitated by the SLC2A family of glucose transporters (GLUTs), encompassing 14 isoforms in humans and 12 homologues in mice (30, 31). GLUT1 has been extensively characterized in conventional T cells, where it is critical for supporting differentiation and effector function (32–36). By contrast, Treg cells appear largely independent of GLUT1, as its deletion had minimal effects on Treg cell homeostasis or suppressive capacity (22, 32), indicating that glucose uptake in Treg cells relies on alternative transporter mechanisms. Our recent studies have revealed that the “neuron type” hexose transporter GLUT3 plays a pivotal role in driving T helper 17 (Th17) effector function through glycolytic-epigenetic reprogramming, highlighting the subset-specific reliance on distinct glucose transporters (11, 37). However, whether GLUT3 similarly regulates the metabolic fitness, epigenetic remodeling and suppressive capacity of Foxp3⁺ Treg cells remains incompletely understood.

In this study, we investigate the role of GLUT3 in Treg physiology using mice with T cell-specific and Treg-intrinsic deletion of GLUT3. We demonstrate that, unlike GLUT1, GLUT3 is essential for the metabolic competence, suppressive capacity and Treg effector differentiation under both steady-state and inflammatory conditions. Inactivation of GLUT3 in Foxp3^+^ Treg cells impairs both glycolytic and mitochondrial activity, leading to fatal autoimmunity driven by diminished Treg numbers and impaired differentiation of effector subsets, including T follicular regulatory (Tfr) cells. These results refine the prevailing notion that Treg cells are largely glucose-independent and identify GLUT3-mediated glucose uptake as a critical regulator of Treg metabolic fitness and peripheral immune tolerance.

## RESULTS

### Ablation of GLUT3 in T cells impairs Treg cell homeostasis

To assess the capacity of Treg cells to uptake and metabolize extracellular glucose, we first analyzed the expression of the GLUT family of hexose transporters in Foxp3^+^ Treg cells and CD4⁺ T conventional (Tcon) cells using publicly available RNA-sequencing datasets (GSE124883) (38). Among the GLUT transporter family, only *Slc2a1* (GLUT1) and *Slc2a3* (GLUT3) were robustly expressed in both Tcon and Treg cells (**Fig. 1A**). Notably, GLUT3 expression was significantly higher in Treg cells compared with Tcon cells. This differential expression was further validated by qPCR in FACS-sorted CD4^+^ Tcon and CD4^+^ CD25^hi^ Treg cells (**Fig. 1B**), supporting a potential role of GLUT3 in Treg cell biology.

**Figure 1.**
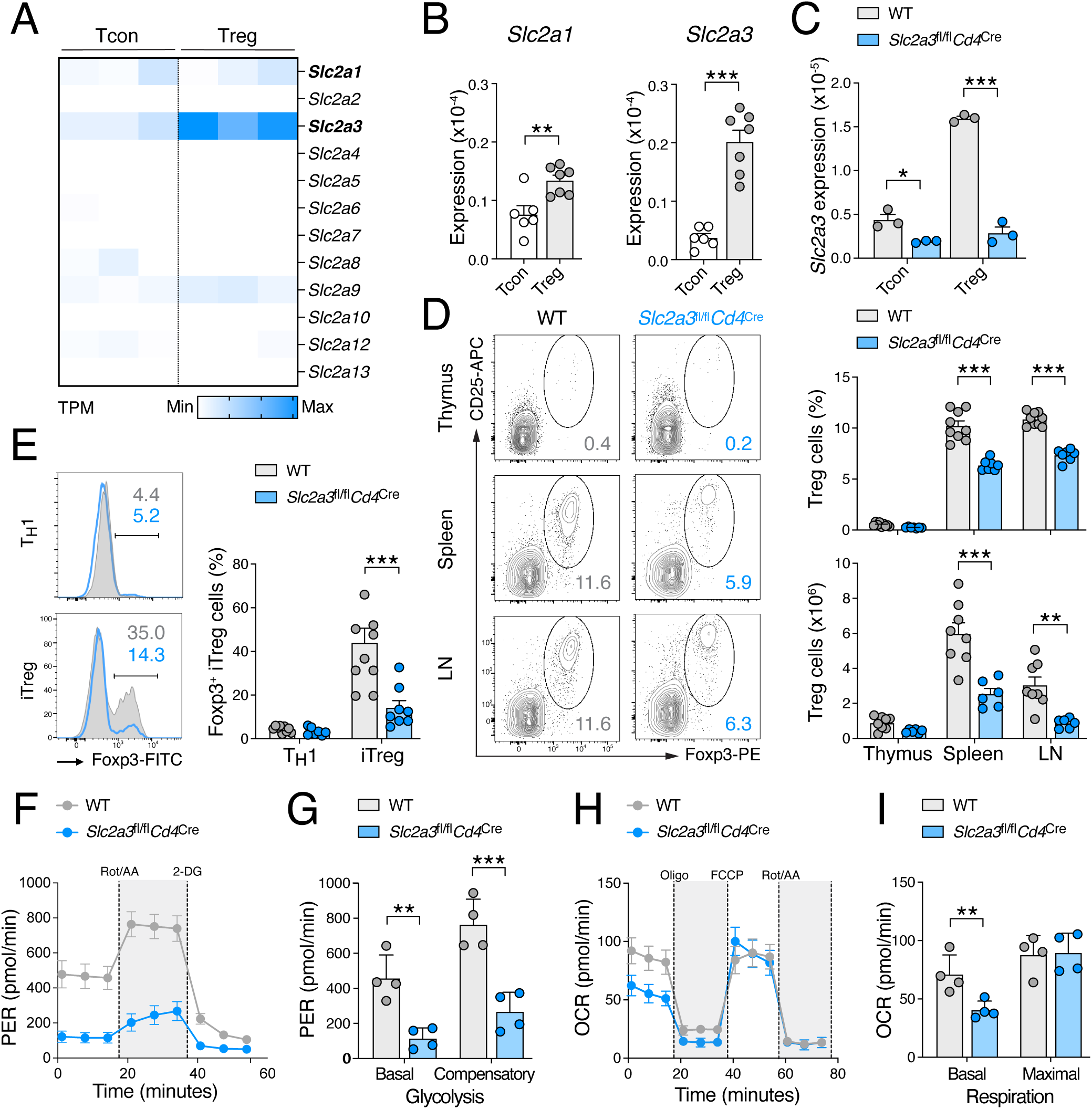
GLUT3 ablation in T cells impairs homeostasis and glycolytic activity of Treg cells. **(A)** Gene expression analysis of GLUT family members (*Slc2a1*–*Slc2a13*) in T conventional (Tcon) and T regulatory (Treg) cells using publicly available RNA-sequencing data (GSE124883); TPM, transcripts per million. **(B)** Expression of *Slc2a1* (GLUT1) and *Slc2a3* (GLUT3) in FACS-sorted *wild type* (WT) Tcon and Treg cells measured by RT-qPCR; data are shown as means ± SEM of 6 mice. **(C)** Analysis of *Slc2a3* gene expression in FACS-sorted Tcon and Treg cells from WT and *Slc2a3*^fl/fl^*Cd4*^Cre^ mice by RT-qPCR; means ± SEM of 3 mice. **(D)** Representative flow cytometric analysis of Foxp3^+^ Treg cells in thymus, spleen and lymph nodes (LNs) of WT and *Slc2a3*^fl/fl^*Cd4*^Cre^ mice (left), with quantification of frequencies and absolute Treg cell numbers (right); means ± SEM of 6-8 mice. **(E)** Analysis Foxp3 expression in *in vitro*-differentiated iTreg cells by flow cytometry; means ± SEM 7-9 mice. **(F** and **G)** Glycolytic proton extrusion rate (PER) in WT and GLUT3-deficient iTreg cells measured using a Seahorse extracellular flux analyzer **(F)**, with quantification of basal and compensatory glycolysis **(G)**; means ± SEM of 4 mice. **(H** and **I)** Analysis of oxygen consumption rate (OCR) in WT and GLUT3-deficient iTreg cells using a Seahorse extracellular flux analyzer **(H)**, with quantification of basal and maximal respiration **(I)**; means ± SEM of 4 mice. Statistical analyses in panels (B-E), (G) and (I) were performed using two-way ANOVA. *, p<0.05; **, p<0.01; ***, p<0.001.

To determine the functional relevance of GLUT3 in Treg cells, we analyzed mice with T cell-specific deletion of GLUT3 (*Slc2a3*^fl/fl^*Cd4*^Cre^ mice) (11) (**Fig. 1C-I**). We first confirmed that *Slc2a3* transcripts were effectively ablated in FACS-sorted CD4^+^ CD25^−^ Tcon and CD4^+^ CD25^hi^ Treg cells from *Slc2a3*^fl/fl^*Cd4*^Cre^ mice compared with *wild type* (WT) controls (**Fig. 1C**). We next examined the frequency and absolute number of CD25^hi^ Foxp3^+^ Treg cells in the thymus, spleen and lymph nodes (LNs) of *Slc2a3*^fl/fl^*Cd4*^Cre^ mice compared to WT littermate controls. Strikingly, T cell-specific ablation of GLUT3 reduced both the relative and absolute cell numbers of Treg cells in the peripheral lymphoid organs by ∼ 30-50 % (**Fig. 1D**). Consistent with this observation, the differentiation of *in vitro*-induced Treg (iTreg) cells from naïve CD4^+^ T cells was markedly impaired in the absence of GLUT3 (**Fig. 1E**). Importantly, we did not observe compensatory upregulation of *Slc2a1* or other GLUT family members in activated GLUT3-deficient T cells (**Fig. S1A**), suggesting that GLUT3 plays a non-redundant role in the differentiation and/or maintenance of Treg cells in the periphery. However, GLUT3-deficient T cells exhibited reduced IL-2 production (**Fig. S1B**), indicating that the impaired generation of iTreg cells may – at least in part – be attributable to limited IL-2 availability.

To further assess the role of GLUT3 in supporting glucose uptake and metabolism in Treg cells, we measured the glycolytic proton efflux rate (PER), an indicator for glycolytic lactate production, and the oxygen consumption rate (OCR), a proxy for mitochondrial respiration, in WT and GLUT3-deficient iTreg cells using a Seahorse extracellular flux analyzer (39). Importantly, both basal and compensatory glycolytic activity were significantly reduced in the absence of GLUT3 (**Fig. 1F,G**), consistent with our previous observations in Th1 and Th17 cells (11). Interestingly, mitochondrial respiration was also markedly impaired in GLUT3-deficient iTreg cells (**Fig. 1H,I**), indicating that GLUT3 ablation not only disrupts aerobic glycolysis but also compromises pyruvate-dependent oxidative metabolism in Foxp3^+^ Treg cells. To determine whether GLUT1 inactivation similarly affects Treg cell biology, we analyzed CD25^hi^ Foxp3^+^ Treg populations in *Slc2a1*^fl/fl^*Cd4*^Cre^ mice (**Fig. S2**). Consistent with previous reports (32), ablation of GLUT1 in T cells did not alter the frequency or absolute cell number of Foxp3^+^ Treg cells in the thymus, spleen and LNs (**Fig. S2A,B**), indicating that GLUT3 fulfills a specific, non-redundant role in Treg cells.

Having established that GLUT3 inactivation impairs Treg cell homeostasis and glycolytic activity at steady state, whereas GLUT1 is dispensable, we next investigated whether these requirements are maintained under inflammatory conditions (**Fig. 2**). We employed experimental autoimmune encephalomyelitis (EAE), a well-established model of T cell-mediated inflammation of the central nervous system (CNS) (40). Although induction and progression of EAE are driven primarily by pathogenic Th1 and Th17 cells, the resolution and termination of neuroinflammation depend on CNS-infiltrating Treg cells (41). We induced EAE in WT, *Slc2a1*^fl/fl^*Cd4*^Cre^ and *Slc2a3*^fl/fl^*Cd4*^Cre^ mice by subcutaneous immunization with MOG_35-55_ peptide emulsified in CFA (40, 41). Consistent with our previous findings (11), *Slc2a3*^fl/fl^*Cd4*^Cre^ mice were almost completely protected from EAE immunopathology, including inflammation-induced weight loss (**Fig. 2A**) and clinical signs of neurological impairment, such as paralysis of the extremities (**Fig. 2B**). By contrast, T cell-specific deletion of GLUT1 did not affect weight loss or disease progression following MOG_35-55_ peptide immunization (**Fig. 2A,B**). 20 days post immunization, we isolated splenic CD4^+^ T cells from *Slc2a1*^fl/fl^*Cd4*^Cre^ and *Slc2a3*^fl/fl^*Cd4*^Cre^ mice and confirmed efficient ablation of GLUT1 and GLUT3, respectively (**Fig. 2C**). The infiltration of CD4^+^ T cells into the spinal cord at the peak of the disease was markedly reduced in *Slc2a3*^fl/fl^*Cd4*^Cre^ mice compared to GLUT1-deficient and control animals (**Fig. 2D,E**). Notably, ablation of GLUT3 also decreased the frequency and absolute number of CNS-resident Treg cells compared with WT and *Slc2a1*^fl/fl^*Cd4*^Cre^ mice (**Fig. 2D,E**). Protection of *Slc2a3*^fl/fl^*Cd4*^Cre^ mice from EAE immunopathology results from defective pathogenic Th17 cell responses (11), as evidenced by the failure of CNS-infiltrating T cells to express GM-CSF (**Fig. 2F,G**) and other pro-inflammatory cytokines. Importantly, analysis of CNS-infiltrating CD4^+^ T cells also revealed a marked reduction in IL-2 production in the absence of GLUT3, whereas GLUT1 ablation did not affect GM-CSF or IL-2 expression (**Fig. 2F,G**). Given the critical role of IL-2 in Treg survival, expansion and tissue recruitment (2, 3), reduced IL-2 availability within the CNS may contribute to the diminished accumulation of Foxp3^+^ Treg cells in the CNS. Because GLUT3-deficient T cells also exhibited impaired IL-2 production *in vitro* (**Fig. S1B**), these findings raise the possibility that defective Treg differentiation and homeostasis under both steady-state and inflammatory condition may be secondary to the reduced IL-2 secretion by conventional T cells.

**Figure 2.**
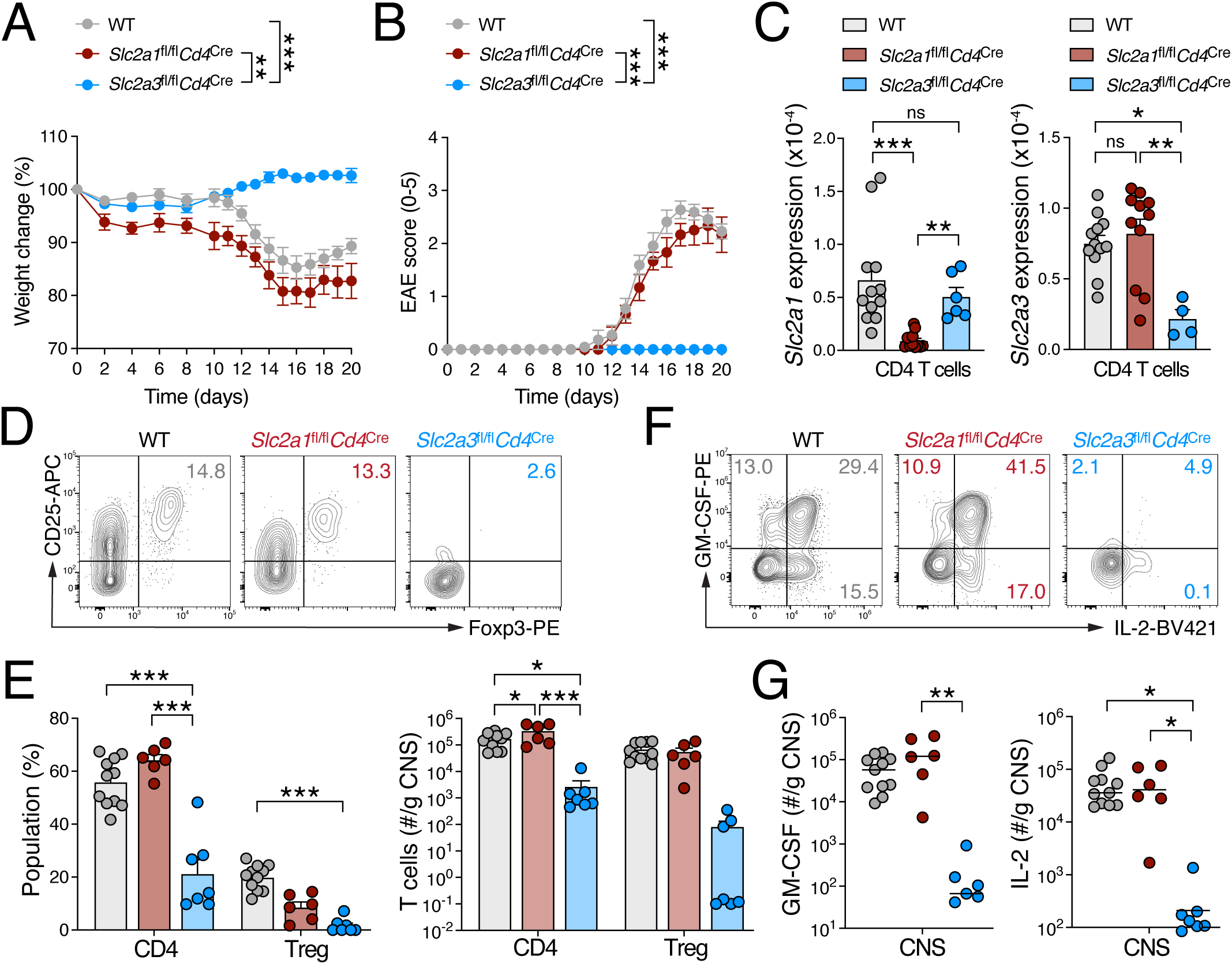
Selective requirement for GLUT3 in sustaining IL-2 production and Treg homeostasis during neuroinflammation. **(A** and **B)** Relative body weight changes **(A)** and experimental autoimmune encephalomyelitis (EAE) clinical scores **(B)** in WT, GLUT1-deficient (*Slc2a1*^fl/fl^*Cd4*^Cre^) and GLUT3-deficient (*Slc2a3*^fl/fl^*Cd4*^Cre^) mice following immunization with MOG_35-55_ peptide emulsified in CFA; means ± SEM of 6-11 mice per cohort. **(C)** RT-qPCR analysis of *Slc2a1* (GLUT1) and *Slc2a3* (GLUT3) expression in isolated splenic CD4^+^ T cells from WT, GLUT1-deficient and GLUT3-deficient mice used in EAE experiments; data are shown as means ± SEM of 4-11 mice. **(D** and **E)** Representative flow cytometric analysis of Foxp3^+^ Treg cells in the central nervous system (CNS) of WT, GLUT1- and GLUT3-deficient mice 20 days after MOG_35-55_ immunization **(D)**, with quantification of Treg cell frequencies and absolute cell numbers **(E)**; means ± SEM of 6-11 mice. **(F** and **G)** Frequencies of GM-CSF and IL-2-producing CD4^+^ T cells in the CNS of WT, GLUT1-and GLUT3-deficient mice 20 days after immunization with MOG_35-55_ peptide and restimulation with PMA/ionomycin for 5 h **(F)**, with quantification of total cell numbers of GM-CSF and IL-2 producing CD4^+^ T cells in the CNS **(G)**; means ± SEM of 6-11 mice. Statistical analyses shown in (A, B, C, E, G) were performed using two-way ANOVA. *, p<0.05; **, p<0.01; ***, p<0.001; ns, non-significant.

### Treg cell-specific ablation of GLUT3 causes early-onset systemic autoimmunity

To clarify whether GLUT3 is intrinsically required for Treg cell metabolic fitness or whether reduced Treg cell numbers result from impaired IL-2 production of conventional T cells, we generated mice with Treg-specific deletion of the *Slc2a3* gene using the *Foxp3*^YFP-Cre^ knock-in mice (42) (hereafter referred to as *Slc2a3*^fl/fl^*Foxp3*^Cre^ mice). Because *Foxp3* is located on the X chromosome, all Foxp3^+^ Treg cells in male hemizygous *Slc2a3*^fl/fl^*Foxp3*^Cre/Y^ mice are deficient for GLUT3, whereas female heterozygous *Slc2a3*^fl/fl^*Foxp3*^Cre/WT^ mice harbor both WT and GLUT3-deficient Treg cells due to random X-chromosome inactivation (4). We first confirmed efficient *Slc2a3* deletion in FACS-sorted CD4^+^ CD25^hi^ Treg cells (**Fig. 3A**). *Slc2a1* (**Fig. 3A**) or other GLUT family members (**Fig. S3A**) were not upregulated in GLUT3-deficient Tregs, arguing against compensation by alternative glucose transporters.

**Figure 3.**
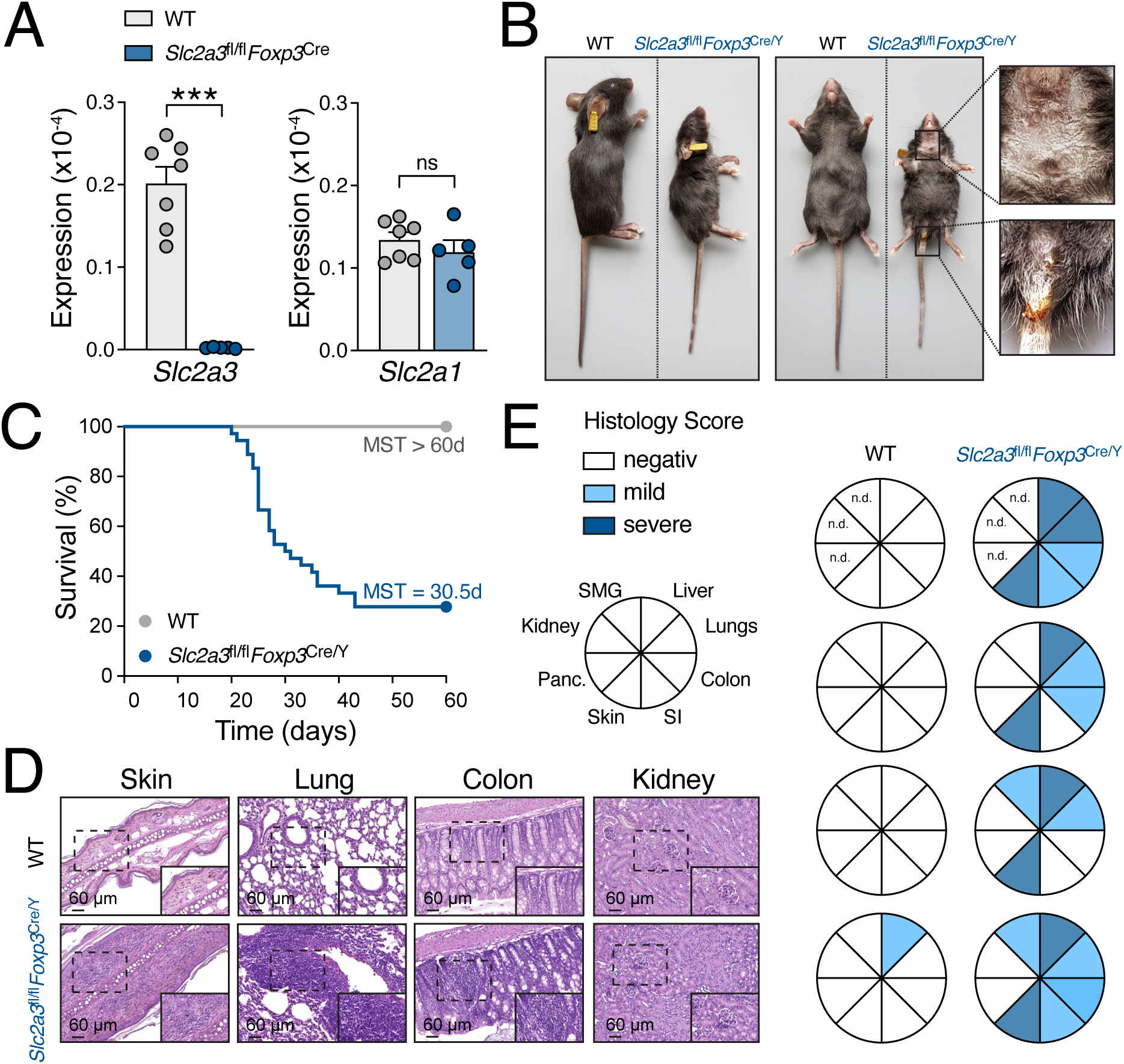
Treg-specific ablation of GLUT3 causes early-onset lethal systemic autoimmunity. **(A)** Analysis of *Slc2a1* (GLUT1) and *Slc2a3* (GLUT3) gene expression in FACS-sorted Treg cells from WT and *Slc2a3*^fl/fl^*Foxp3*^Cre^ mice by RT-qPCR; data are shown as means ± SEM of 5-7 mice. **(B)** Representative images of 28-days old hemizygous male WT and *Slc2a3*^fl/fl^*Foxp3*^Cre/Y^ littermate mice. **(C)** Kaplan-Meier survival curves of hemizygous male WT and *Slc2a3*^fl/fl^*Foxp3*^Cre/Y^ mice; >35 mice per cohort. Mean survival time (MST) was 30.5 days for *Slc2a3*^fl/fl^*Foxp3*^Cre/Y^ mice and > 60 days for WT mice. **(D)** Representative examples of hematoxylin and eosin (H&E)-stained sections of skin, lung, colon and kidney from male WT and *Slc2a3*^fl/fl^*Foxp3*^Cre/Y^ mice; scale bars represent 60 µm. **(E)** Histopathological assessment of leukocytic infiltration and tissue inflammation in non-lymphoid organs from hemizygous male WT and *Slc2a3*^fl/fl^*Foxp3*^Cre/Y^ mice by histopathology, as shown in (D); each pie chart represents one individual mouse. Statistical analyses in panel (A) were performed using unpaired Student’s t-tests. ***, p<0.001; ns, non-significant.

Strikingly, male *Slc2a3*^fl/fl^*Foxp3*^Cre/Y^ mice developed a rapidly progressing inflammatory syndrome resembling the “scurfy” phenotype observed in mice lacking functional Foxp3 expression (1, 4–7) (**Fig. 3B**). *Slc2a3*^fl/fl^*Foxp3*^Cre/Y^ animals were born at the expected Mendelian ratio but exhibited growth retardation, ruffled fur, scurfy skin lesions and signs of diarrhea (**Fig. 3B**). Consequently, 50 % of the affected *Slc2a3*^fl/fl^*Foxp3*^Cre/Y^ mice succumbed to disease by ∼ 30 days *postpartum* (**Fig. 3C**), demonstrating a critical role for GLUT3 in Treg cells. Histological examination of multiple barrier and non-barrier tissues from *Slc2a3*^fl/fl^*Foxp3*^Cre/Y^ and littermate controls revealed extensive leukocytic infiltration and marked tissue inflammation in the skin, lungs and liver, whereas other organs, such as the kidneys, submandibular glands (SMG), colon, small intestine and pancreas were less affected (**Fig. 3D,E** and **Fig. S3B**). Together, these findings show that GLUT3 is intrinsically required in Foxp3^+^ Treg cells to prevent widespread tissue inflammation, resulting in fatal systemic autoimmunity.

Inspection of lymphoid organs in *Slc2a3*^fl/fl^*Foxp3*^Cre/Y^ mice revealed pronounced lymphadenopathy, with significantly enlarged submandibular, inguinal and axillary LNs compared to WT littermate controls, whereas mesenteric LNs were less affected (**Fig. 4A,B**). Flow cytometric analysis of CD4^+^ and CD8^+^ T cell compartments in spleen and LNs further demonstrated a marked shift from naïve to effector/memory phenotype (**Fig. 4C,D** and **Fig. S3C,D**). Given this broad immune activation, we next assessed cytokine profiles in the sera of male WT and *Slc2a3*^fl/fl^*Foxp3*^Cre/Y^ littermate mice to characterize the inflammatory syndrome. Serum concentrations of IFNγ and TNFα were significantly elevated, consistent with a type I inflammatory response (**Fig. 4E**). In addition, cytokines associated with type II immunity, including IL-4, IL-5 and IL-13, were markedly increased, with IL-5 elevated by ∼ 130-fold in the sera of *Slc2a3*^fl/fl^*Foxp3*^Cre/Y^ mice compared to WT littermates (**Fig. 4E**). Correspondingly, the frequency of CD11b^+^ Siglec F^+^ eosinophiles was markedly increased in both lymphoid tissues and non-lymphoid organs such as the lung (**Fig. 4F**).

**Figure 4.**
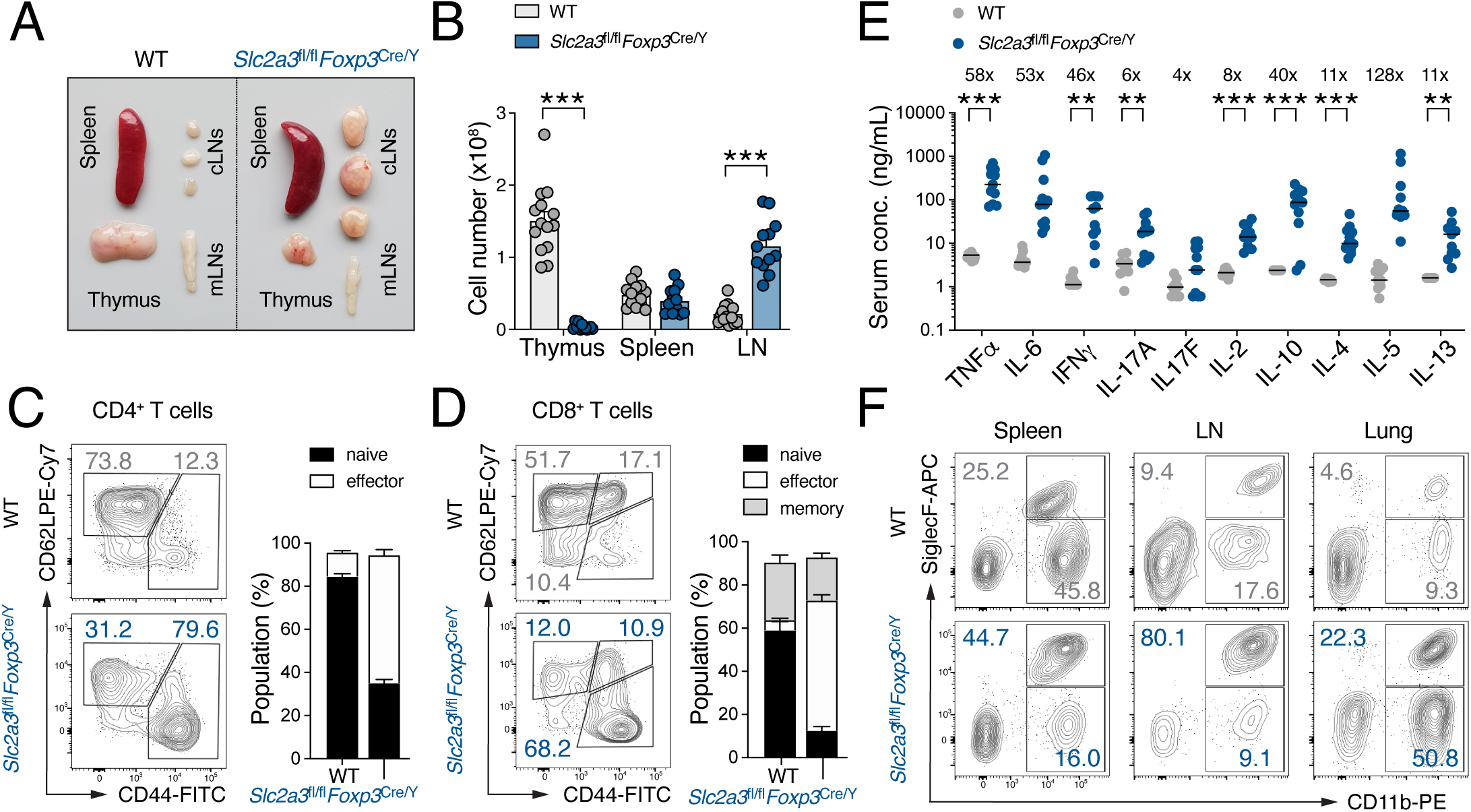
Systemic immune cell activation in mice with Treg-specific GLUT3 ablation. **(A)** Macroscopic images of thymus, spleen, cervical and mesenteric lymph nodes (LNs) from male WT and hemizygous *Slc2a3*^fl/fl^*Foxp3*^Cre/Y^ mice. **(B)** Total cell numbers in thymus, spleen and LNs of WT and *Slc2a3*^fl/fl^*Foxp3*^Cre/Y^ mice; data are shown as means ± SEM of 12-14 mice. **(C** and **D)** Flow cytometric analysis of naïve (CD62L^hi^ CD44^lo^), effector (CD62L^lo^ CD44^hi^) and central memory (CD62L^hi^ CD44^hi^) CD4^+^ **(C)** and CD8^+^ **(D)** T cells in LNs of male WT and *Slc2a3*^fl/fl^*Foxp3*^Cre/Y^ mice; means ± SEM of 10-13 mice. **(E)** Serum cytokine levels in male WT and hemizygous *Slc2a3*^fl/fl^*Foxp3*^Cre/Y^ mice; fold increase in *Slc2a3*^fl/fl^*Foxp3*^Cre/Y^ mice relative to littermate WT mice are indicated by “x”; means ± SEM of 9-11 mice. **(F)** Representative flow cytometric analysis of eosinophils in the spleen, LNs and lung tissue of male WT and *Slc2a3*^fl/fl^*Foxp3*^Cre/Y^ mice. Statistical analyses in panels (B-E) were performed using two-way ANOVA. *, p<0.05; **, p<0.01; ***, p<0.001.

Collectively, these results demonstrate that GLUT3 is intrinsically required in Treg cells to restrain systemic immune activation, eosinophilia and fatal autoimmunity in male mice.

### GLUT3 regulates effector differentiation of Foxp3^+^ Treg cells

To further characterize the impact of GLUT3 deficiency on Foxp3^+^ Treg cells, we analyzed the abundance and phenotype of central and effector Treg cell subsets in *Slc2a3*^fl/fl^*Foxp3*^Cre/Y^ mice (**Fig. 5**). Consistent with the observations in mice lacking GLUT3 in all T cells (**Fig. 1D**), Treg cell-specific deletion of GLUT3 reduced the frequency of Foxp3^+^ Treg cells within the CD4^+^ T cell compartment in the spleen and LNs (**Fig. 5A**). However, despite this decrease in relative frequency, the absolute number of Treg cells in the LNs was increased due to the overall elevated cellularity (**Fig. S4A**).

**Figure 5.**
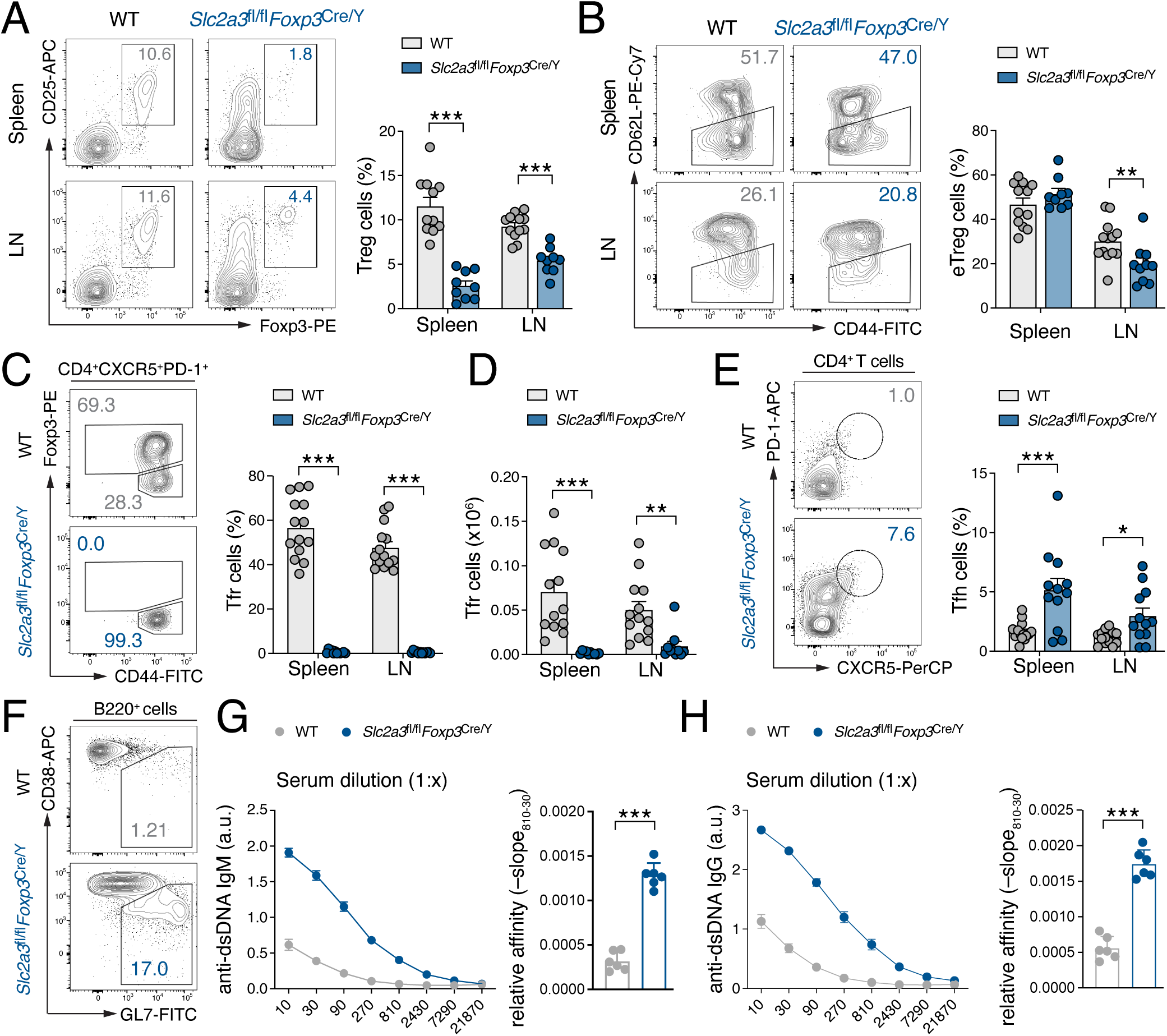
GLUT3 controls Treg cell homeostasis and effector differentiation. **(A** and **B)** Representative flow cytometric analysis of Foxp3^+^ Treg cells in spleen and lymph nodes (LNs) of male WT and hemizygous *Slc2a3*^fl/fl^*Foxp3*^Cre/Y^ mice (left), with quantification of Treg cell frequencies (right); data are shown as means ± SEM of 10-13 mice. **(B)** Analysis of central (CD62L^hi^ CD44^hi^) and effector (CD62L^lo^ CD44^hi^) Treg cells in spleen and LNs of male WT and *Slc2a3*^fl/fl^*Foxp3*^Cre/Y^ mice by flow cytometry; means ± SEM of 10-13 mice. **(C** and **D)** Flow cytometric analysis of Foxp3^+^ T follicular regulatory (Tfr) cells in the spleen and LNs of male WT and *Slc2a3*^fl/fl^*Foxp3*^Cre/Y^ mice, with quantification of Tfr cell frequencies **(C)** and absolute cell numbers **(D)**; means ± SEM of 12-14 mice. **(E)** Flow cytometric analysis of CD4^+^ T follicular helper (Tfr) cells in the spleen and LNs of male WT and *Slc2a3*^fl/fl^*Foxp3*^Cre/Y^ mice, with quantification of Tfh cell frequencies, means ± SEM of 12-14 mice. **(F)** Representative flow cytometric analysis of CD38^lo^ GL7^hi^ germinal center (GC) B cells in LNs of male WT and *Slc2a3*^fl/fl^*Foxp3*^Cre/Y^ mice. **(G** and **H)** ELISA quantification of anti-double-stranded DNA (dsDNA) IgM **(G)** and IgG **(H)** autoantibody titers and relative affinity in sera of hemizygous male WT and *Slc2a3*^fl/fl^*Foxp3*^Cre/Y^ littermate mice; means ± SEM of 6 mice. Statistical analyses in panels (A-E) were performed using two-way ANOVA. Statistical analysis shown in (G and H) by unpaired Student’s t-tests. *, p<0.05; **, p<0.01, ***, p<0.001.

Thymus-derived Treg cells can be broadly divided into circulating CD62L^+^ CD44^lo^ central Treg (cTreg) cells and CD62L^−^ CD44^hi^ effector Treg (eTreg) cells (43–46). Notably, the eTreg cell subset was markedly reduced in the LNs of *Slc2a3*^fl/fl^*Foxp3*^Cre/Y^ mice compared to WT littermates (**Fig. 5B** and **Fig. S4B**). Among the eTreg population, T follicular regulatory (Tfr) cells represent a specialized subset that controls germinal center (GC) responses. Tfr cells arise from Foxp3⁺ Treg cells and acquire expression of CXCR5 and the transcription factor Bcl6, enabling their migration into B cell follicles (46). Within B cell follicles, Tfr cells restrain T follicular helper (Tfh) cells and regulate GC responses, thereby limiting excessive antibody responses and maintaining humoral immune tolerance (46–49). Strikingly, GLUT3-deficient Treg cells were unable to differentiate into CD44^+^ CXCR5^+^ PD-1^+^ Foxp3^+^ Tfr cells (**Fig. 5C,D**), resulting in the accumulation of CD4^+^ CXCR5^+^ PD-1^+^ Tfh cell (**Fig. 5E**) and spontaneous formation of B220^+^ CD38^−^ GL7^+^ GC B cells (**Fig. 5F** and **Fig. S4C,D**). Consequently, male *Slc2a3*^fl/fl^*Foxp3*^Cre/Y^ mice exhibited pathologically elevated serum levels of anti-dsDNA IgM and IgG autoantibodies compared with WT littermates (**Fig. 5G,H**).

By contrast, none of these effects were observed in mice with Treg cell-specific ablation of GLUT1 (*Slc2a1*^fl/fl^*Foxp3*^Cre/Y^ mice) (**Fig. S5A-F**), demonstrating that GLUT1 and GLUT3 play non-redundant roles in eTreg cell differentiation and in restraining cellular and humoral autoimmunity.

### GLUT3 is required for the metabolic fitness and suppressive capacity of Treg cells

To further investigate the cell-intrinsic role of GLUT3 in Treg cells under non-inflammatory conditions, we took advantage of female heterozygous *Slc2a3*^fl/fl^*Foxp3*^Cre/WT^ mice. In contrast to male hemizygous *Slc2a3*^fl/fl^*Foxp3*^Cre/Y^ mice, in which all Treg cells lack GLUT3, female heterozygous mice harbor both WT and GLUT3-deficient Treg cells within the same animal due to lyonization (that is, random X chromosome inactivation) (**Fig. 6A**). The presence of Foxp3^+^ YFP^−^ WT Treg cells in these females is sufficient to maintain normal immune tolerance despite the coexistence of potentially dysfunctional GLUT3-deficient Foxp3^+^ YFP^+^ Treg cells (**Fig. 6B**), making this model an ideal system to study the role of GLUT3 in Treg cells without the confounding effects of overt immune dysregulation.

**Figure 6.**
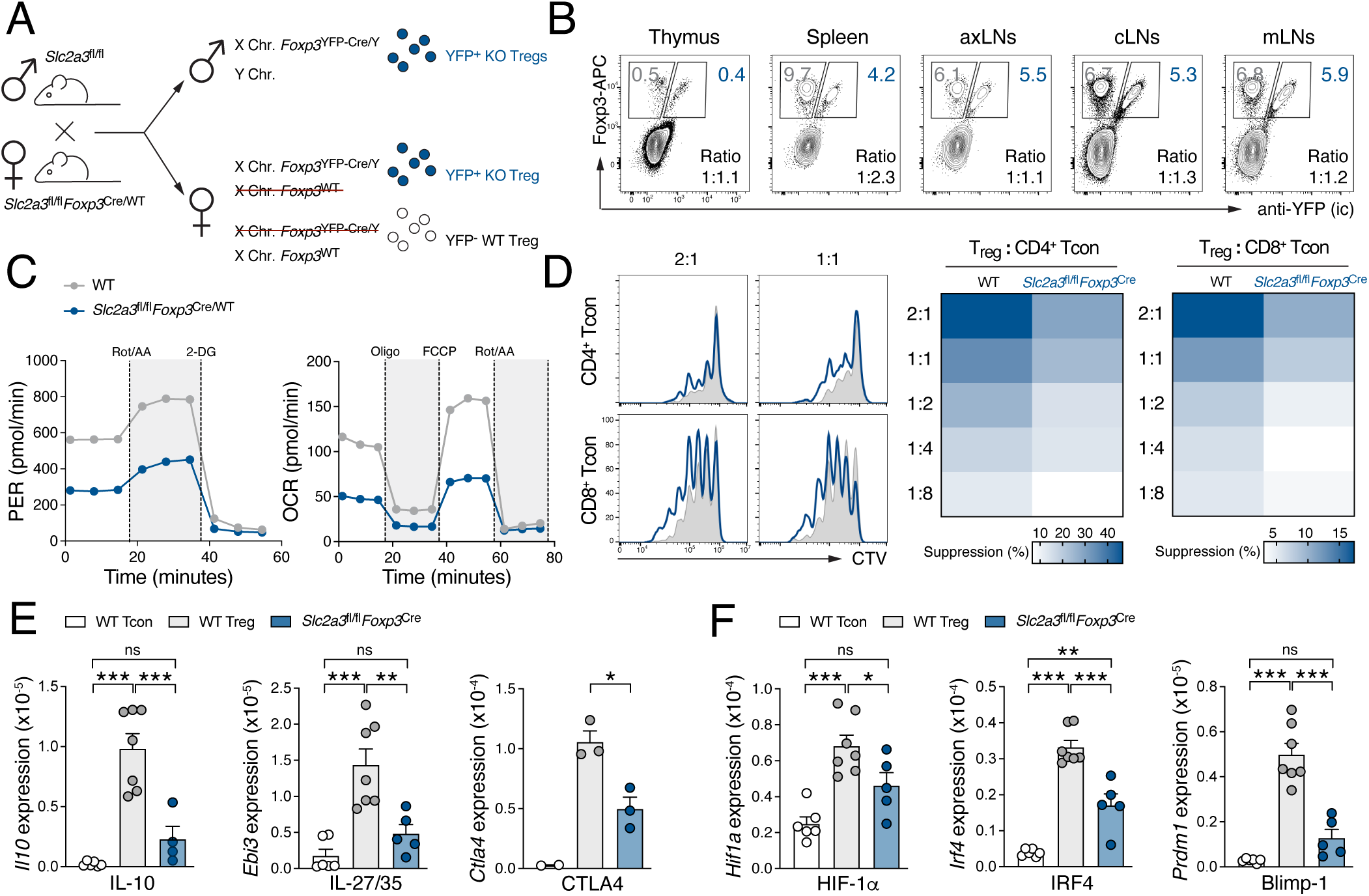
GLUT3 is intrinsically required in Treg cells for aerobic glycolysis, suppressive function and effector differentiation. **(A)** Female heterozygous *Slc2a3*^fl/fl^*Foxp3*^Cre/+^ mice harbor both GLUT3-sufficient (negative for *Foxp3*^Cre^) and GLUT3-deficient (express *Foxp3*^Cre^) Treg cells in the same animal due to random X chromosome inactivation. Male hemizygous *Slc2a3*^fl/fl^*Foxp3*^Cre/Y^ mice, in which all Treg cells are GLUT3-deficient, develop systemic autoimmunity, whereas female heterozygotes remain healthy due to the presence of WT Treg cells. **(B)** Representative flow cytometric analysis of intracellular Foxp3 and YFP protein expression in WT and GLUT3-deficient Treg cells from different lymphoid organs of female heterozygous *Slc2a3*^fl/fl^*Foxp3*^Cre/+^ mice. **(C)** Seahorse extracellular flux analyses of glycolytic proton extrusion rate (PER) and oxygen consumption rate (OCR) in WT and GLUT3-deficient Treg cells isolated from female heterozygous *Slc2a3*^fl/fl^*Foxp3*^Cre/+^ mice; data are representative of 5 pooled mice. **(D)** Suppression of T cell proliferation by WT and GLUT3-deficient Treg cells. CellTrace Violet (CTV)-labeled responder T cells were co-cultured with varying ratios of WT and GLUT3-deficient Treg cells isolated from female heterozygous *Slc2a3*^fl/fl^*Foxp3*^Cre/+^ mice. CTV dilution of CD4^+^ and CD8^+^ responder T cell was analyzed by flow cytometry; data are representative of two independent experiments. **(E** and **F)** Expression of selected suppressive molecules **(E)** and transcription factors promoting effector Treg cell differentiation **(F)** in FACS-sorted WT and GLUT3-deficient Treg cells from female heterozygous *Slc2a3*^fl/fl^*Foxp3*^Cre/+^ mice measured by RT-qPCR; data are shown as means ± SEM of 3-7 mice. Statistical analyses in panels (E) and (F) were performed using two-way ANOVA. *, p<0.05; **, p<0.01; ***, p<0.001; ns, non-significant.

We next isolated WT and GLUT3-deficient Foxp3^+^ Treg cells from *Slc2a3*^fl/fl^*Foxp3*^Cre/WT^ mice by FACS-sorting (**Fig. S6A**) and assessed the glycolytic activity (PER) and mitochondrial respiration (OCR) using a Seahorse extracellular flux analyzer (**Fig. 6C**). Both glycolytic flux and oxygen consumption were reduced by ∼ 50 % in GLUT3-deficient Treg cells compared to WT controls from the same animal, similar to the reduction observed in *Slc2a3*^fl/fl^*Cd4*^Cre^ mice (**Fig. 1F-I**). To further determine the suppressive capacity of WT and GLUT3-deficient Treg cells *in vitro*, FACS-sorted WT and GLUT3-deficient Treg cells were co-cultured with CellTrace Violet (CTV)-labeled CD45.1^+^ T cells in different Treg-to-Tcon ratios. Although GLUT3-deficient Treg cells retained some suppression toward the proliferation of activated CD4^+^ and CD8^+^ T cells, their suppressive capacity was markedly reduced compared with WT Treg cells (**Fig. 6D**). This functional impairment was associated with decreased expression of key immunoregulatory molecules, including *Il10* (IL-10), *Ebi3* (IL-35), *Gzmb* (granzyme B) and *Ctla4* (CTLA-4) (**Fig. 6E** and **Fig. S6B**). Given that GLUT3-deficient Treg cells failed to differentiate into Tfr cells (**Fig. 5C,D**), we next examined selected transcription factors known to regulate the effector differentiation of Treg cells (15, 26, 44, 46). Expression of *Hif1a* (HIF-1a), *Irf4* (IRF-4), *Prdm1* (Blimp-1) and *Gata3* (GATA-3) was markedly reduced in GLUT3-deficient Treg cells, whereas expression of *Tcf7* (TCF-1), which also regulates eTreg differentiation (50), remained unchanged compared with WT controls (**Fig. 6F** and **Fig. S6C**).

Altogether, these results demonstrate that GLUT3 plays a critical cell-intrinsic role in maintaining the metabolic fitness, suppressive function and effector differentiation of Foxp3^+^ Treg cells, thereby preserving normal immune homeostasis.

## DISCUSSION

Treg cells rely on tightly coordinated metabolic programs to sustain their differentiation, lineage stability, and suppressive function (51, 52). Over the past decade, immunometabolism has emerged as a central determinant of T cell fate, revealing that metabolic pathways actively instruct, rather than merely support, immune cell identity and function (53–55). Unlike conventional effector T cells, which strongly induce aerobic glycolysis after antigen receptor stimulation to fuel proliferation and cytokine production, Treg cells have traditionally been considered to depend primarily on mitochondrial OxPhos and fatty acid oxidation to maintain their suppressive program. This oxidative state has been associated with Treg longevity, functional stability and persistence *in vivo* (55). Accordingly, disruption of mitochondrial integrity, electron transport chain (ETC) activity or mitochondrial regulatory pathways impairs Treg suppressive function and destabilizes Foxp3 expression, thereby compromising immune homeostasis (21, 53, 54). In addition to ATP generation, mitochondrial metabolism supplies biosynthetic intermediates that shape chromatin remodeling and epigenetic programs required for Treg identity and maintenance (21, 56). Moreover, tricarboxylic acid (TCA) cycle intermediates and physiological reactive oxygen species (ROS) generated by the ETC complexes act as key signaling mediators linking metabolic activity to intracellular signaling pathways and Foxp3 stability (28, 57–61).

Despite their heightened mitochondrial capacity and reliance on oxidative metabolism, accumulating evidence indicates that Foxp3⁺ Treg cells retain substantial metabolic flexibility and can engage aerobic glycolysis in a context-dependent manner (20, 25, 29, 62). Glycolytic metabolism has been implicated in Treg proliferation, migration and the generation of effector Treg cells, as well as in the acquisition of tissue-adapted phenotypes, suggesting that distinct Treg subsets rely on different metabolic modules depending on the anatomical location, metabolic environment and inflammatory niche (20, 25, 29, 60, 62, 63). Antigen receptor signaling, co-stimulation, IL-2 responsiveness, hypoxia and environmental nutrient cues have emerged as important determinants of Treg metabolic state and adaptability (64). These inputs converge on central nutrient-sensing pathways, including mTOR, AMPK and PI3K-Akt, which integrate extracellular signals with intracellular metabolic programming to balance quiescence, proliferation, lineage stability and suppressive function (20, 52, 65). An additional layer of control is provided by Foxp3 itself, which not only orchestrates the canonical immunoregulatory transcriptional program of Treg cells but also contributes directly to their metabolic phenotype. Foxp3 has been shown to adjust the expression of glucose transporters and glycolytic enzymes following T cell receptor stimulation, thereby limiting excessive glycolytic flux and promoting a metabolic state compatible with suppressive activity (18, 22). Conversely, loss of Foxp3 results in excessive glycolytic activity accompanied by impaired Treg stability and diminished immunoregulatory capacity (23). Together, these findings support a model in which Treg cell identity is maintained by a finely tuned balance between mitochondrial and glycolytic metabolism, rather than by absolute commitment to a single bioenergetic pathway.

The glycolytic program of Treg cells appears to be particularly important during activation, migration, and effector differentiation (20, 25, 29, 62). Notably, glycolysis-dependent mTOR signaling drives cytoskeletal remodeling required for Treg cell migration into inflamed tissues (25, 62). Moreover, glycolytic flux and HIF-1α signaling can directly modulate Foxp3 stability, whereas glucose-derived metabolites may serve as substrates for post-translational modifications that influence Treg-mediated antitumor immunity (28, 60). A conserved role for glucose metabolism is further supported by findings in human Treg cells, in which TNF receptor signaling induces a glycolytic switch associated with enhanced proliferation and effector function (63). Given that hexose transporter expression defines cellular glucose uptake capacity, we here investigated the role of the two principal glucose transporters, GLUT1 and GLUT3, in Foxp3⁺ Treg cell physiology. In conventional effector T cells, GLUT1 has emerged as the dominant glucose transporter that supports activation, proliferation and effector differentiation, consistent with their strong dependence on aerobic glycolysis (32, 35). By contrast, the relatively modest impact of GLUT1 deficiency on Treg homeostasis and suppressive capacity (22, 32) suggest that Treg cells rely on alternative glucose transport mechanisms. In this regard, our previous work identified the “neuron type” glucose transporter GLUT3 as a driver of glucose-dependent acetyl-CoA generation and metabolic-epigenetic remodeling in effector T cells (11). Extending this concept to the Treg lineage, we here demonstrate that GLUT3, but not GLUT1, is essential for Foxp3⁺ Treg cell homeostasis and functional specification.

Although Tregs are metabolically flexible, our findings identify a critical requirement for GLUT3-mediated glucose uptake in maintaining their metabolic fitness, suppressive function and effector differentiation. GLUT3-dependent glucose availability was required to sustain both glycolytic and mitochondrial activity, particularly in effector Treg subsets, underscoring the importance of glucose metabolism in these cells. Using both T cell-specific and Treg-specific GLUT3 ablation models, we demonstrate that this requirement is cell intrinsic and essential for the prevention of lethal systemic inflammation. GLUT3 deficiency in Treg cells led to a marked reduction in overall Treg abundance in lymphoid and non-lymphoid tissues, diminished suppressive capacity and profound defects in functional specialization, such as Tfr cell differentiation. Accordingly, mice lacking GLUT3 in Tregs developed a severe early-onset autoimmune syndrome marked by multiorgan leukocytic infiltration, spontaneous GC formation and pathogenic autoantibody production. Notably, this phenotype was not reproduced by GLUT1 deficiency, indicating that GLUT3 fulfills a non-redundant role and emphasizing that different glucose transporters have distinct contributions in lymphocyte biology.

A recent report showed that tumor-infiltrating Treg cells upregulate the high-affinity glucose transporter GLUT3 in response to the glucose-deprived tumor microenvironment (28, 66). Using mice with Treg-specific GLUT3 deficiency, Sharma *et al*. found that loss of GLUT3 selectively reduced the abundance and suppressive capacity of tumor-infiltrating Treg cells, while having only limited effects on other Treg cell subsets and on systemic immune tolerance. These findings contrast with our results, in which Treg-specific GLUT3 ablation caused systemic immune activation and severe immunopathology. One possible explanation for this discrepancy is the difference in genetic targeting strategies; Sharma *et al.* deleted exons 2 and 3 of the *Slc2a3* gene locus, whereas our model is based on conditional deletion of exon 7, which may have contributed to the divergent phenotypic outcomes. Despite these differences in phenotype, both studies support the conclusion that GLUT3 is required to sustain Treg cell metabolism, with loss of GLUT3 leading to impaired glycolysis and reduces the availability of glucose-derived pyruvate to support mitochondrial respiration. However, the functional consequences of defective glucose uptake are likely not limited to cellular bioenergetics. Glucose can also be redirected into the biosynthesis of unsaturated fatty acids that promote Treg proliferation and suppressive function (63). In addition, GLUT3-dependent glucose metabolism may influence post-translational protein modifications, including acetylation, lactylation, methylation and other metabolite-dependent modifications linked to glycolytic and TCA cycle intermediates (11, 21, 28). Consistent with this notion, Sharma *et al.* identified GLUT3-dependent O-GlcNAcylation of the transcription factor c-Rel as a critical regulatory mechanism in tumor-infiltrating Treg cells (28).

Altogether, our findings establish GLUT3-dependent glucose uptake as a critical regulator of the metabolic fitness and functional specification of Foxp3^+^ Treg cells, thereby maintaining cellular and humoral immune tolerance. Our results also have important clinical implications, as targeting GLUT1/3-dependent immunometabolism has emerged as a promising strategy for treating inflammatory and autoimmune diseases (11, 67–69). Given the pivotal role of Treg cells in maintaining immune balance and restraining inflammation, targeting glucose uptake in both effector and regulatory T cells may inadvertently exacerbate immune dysregulation.

## DATA LIMITATION AND PERSPECTIVES

Treg cell-specific deletion of GLUT3 resulted in an early-onset, fatal autoimmune syndrome, a phenotype that was not observed in mice with GLUT3 ablation across all T cells. We attribute this discrepancy to the requirement of GLUT3 in conventional T cells, including Th1 and Th17 subsets, for their pathogenic effector functions (11). Thus, the simultaneous loss of GLUT3 in both effector T cells and Treg cells may lead to a functional balance between inflammatory and regulatory responses. However, *Slc2a3*^fl/fl^*Cd4*^Cre^ mice exhibited an increased incidence of prolapse with age, suggesting that Treg cell-mediated immunoregulation is also compromised in these mice. It is noteworthy that not all male hemizygous *Slc2a3*^fl/fl^*Foxp3*^Cre/Y^ mice developed fatal autoimmune disease, as shown in Fig. 3C. Analysis of GLUT3 expression in FACS-sorted Treg cells by RT-qPCR indicated incomplete deletion in these animals, which may account for this variability. Finally, although no overt alterations in Treg cell number and phenotype were observed in mice with Treg-intrinsic GLUT1 deletion, these findings do not preclude a context-dependent requirement for GLUT1 in Treg cells under specific physiological or pathological conditions, such as tissue-specific inflammation or in the microenvironment of tumors.

## MATERIALS AND METHODS

### Mice

All mice were bred and maintained under specific pathogen-free conditions at the Center for Experimental Medicine (ZEMM) or the Institute for Systems Immunology at the Julius-Maximilians-University of Würzburg, Germany. Animals were housed in individually ventilated cages under a 12 h light/12 h dark cycle at 20-24 °C. Mice had *ad libitum* access to standard chow and autoclaved drinking water. The hygienic status of sentinel mice was monitored quarterly in accordance with FELASA guidelines. Male hemizygous *Slc2a3*^fl/fl^*Foxp3*^Cre/Y^ mice were used at ∼ 4 weeks of age, whereas female heterozygous *Slc2a3*^fl/fl^*Foxp3*^Cre/+^ mice were used between 8-24 weeks of age. WT littermates served as age- and sex-matched controls. All animal procedures were approved by the Government of Lower Franconia, Germany. C57BL/6 (stock no. 000664), *Cd4*^Cre^ (017336), *Foxp3*^Cre^ (016959) and *Slc2a1*^fl/fl^ (031871) mice were purchased from the Jackson Laboratories (JAX) and maintained in our institution. *Slc2a3*^fl/fl^ mice were generated through the Knockout Mouse Project (KOMP) at UC Davis (stock number 049702-UCD) and have been previously described (11, 70). All animals used in this study were maintained on a C57BL/6 genetic background.

### Experimental autoimmune encephalomyelitis (EAE)

EAE was induced by emulsifying 2 mg/mL myelin MOG_35–55_ peptide (Synpeptide) in incomplete Freund’s adjuvant (IFA; Fisher Scientific) supplemented with 5 mg/mL *Mycobacterium tuberculosis* H37Ra (Fisher Scientific) via syringe extrusion. Each mouse received 200 μg MOG_35–55_ peptide injected subcutaneously (s.c.) at two sites on the lower flanks, followed by intraperitoneal (i.p.) administration of 250 ng pertussis toxin (Enzo Life Sciences) on days 0 and 2. Mice were monitored daily for clinical signs of EAE and body weight changes. Clinical scores were assigned as previously described (40): 0, no paralysis; 0.5, partially limp tail; 1, paralyzed tail; 2, uncoordinated movement; 2.5, paralysis of one hind limb; 3, paralysis of both hind limbs; 3.5, forelimb weakness; 4, forelimb paralysis; 5, moribund. At the experimental endpoint, mice were sacrificed, and single-cell suspensions were prepared from spleen, LNs and spinal cords. For CNS isolation, spinal cords were excised, minced into small fragments and digested with a cocktail of 1 mg/mL collagenase D (Roche) and 20 μg/mL DNase I (Sigma-Aldrich) for 40 min at 37 °C. After filtration through a 70 μm cell strainer, CNS-infiltrating lymphocytes were enriched by Percoll (Sigma-Aldrich) gradient centrifugation (30:70) and analyzed by flow cytometry.

### Flow cytometry and cell sorting

Flow cytometric staining was performed as previously described (49). Briefly, cells were stained with Fixable Viability Dye eFluor 780 (eBioscience) for 10 min in PBS at RT together with an anti-CD16/CD32 antibody (clone 2.4G2; Bio X Cell) to prevent non-specific binding. After washing, surface antigens were detected with fluorophore-conjugated antibodies (**Table S1**) in PBS containing 0.5 % BSA for 20 min at RT in the dark. For the analysis of intranuclear antigens, cells were fixed using the Foxp3/Transcription Factor staining buffer set (eBioscience) according to manufacturer’s instructions and subsequently stained with antibodies against transcription factors in 1x permeabilization buffer for 40 min at RT (eBioscience). For intracellular cytokine staining, cells were stimulated with 1 μM ionomycin (BioMol) and 30 nM phorbol-12-myristat-13-acetate (PMA; Sigma-Aldrich) in the presence of 3 μg/ml brefeldin A and 2 μM monensin (both eBioscience) for 4-5 h at 37 °C. After surface staining, cells were fixed with IC Fixation Buffer (eBioscience) and intracellular cytokines were stained using 1x permeabilization buffer (eBioscience) for 40 min at RT. All samples were acquired on a BD FACSCelesta flow cytometer (BD Biosciences) or an Aurora Flow Cytometer (Cytek Biosciences) and further analyzed with the FlowJo software (BD Biosciences). For the isolation of Tcon and Treg cells by FACS sorting, total CD4^+^ T cells were enriched using the MojoSort CD4 T Cell Isolation Kit (BioLegend). After surface staining, the respective cell populations were sorted using an BD FACSAria III Cell Sorter.

### T cell cultivation and differentiation

For *in vitro* cultures, murine CD4^+^ T cells were isolated from single-cell suspension of lymph nodes and spleen by negative selection using the MojoSort Mouse CD4^+^ T Cell Isolation Kit (BioLegend). T cells were cultured in modified RPMI 1640 medium containing a physiological glucose concentration (100 mg/dL) by diluting standard RPMI 1640 medium (Gibco) with glucose-free RPMI medium (Carl Roth). The medium was supplemented with 10 % FBS (Sigma-Aldrich), 50 μM 2-mercaptoethanol, 1 % penicillin-streptomycin and 1 % GlutaMAX (all Gibco). For T cell differentiation and activation, Delta surface culture plates (Nunc) were pre-coated with 12 μg/mL polyclonal anti-hamster IgG (MP Biomedicals) for at least 2 h and washed once with PBS. In 24-well plates, 1×10^6^ cells were activated with plate-bound antibodies consisting of 0.5 μg/mL anti-CD3 (clone 145-2C1) and 1 μg/mL anti-CD28 (clone 37.51) and polarized into different T cell subsets with the following cytokine and antibody combinations. For Th1 differentiation, cells were cultured with 2.5 μg/mL anti-IL-4 (clone 11B11, Bio X Cell), 10 ng/mL recombinant human IL-2 and 10 ng/mL recombinant murine IL-12 (both PeproTech). For iTreg differentiation, CD4^+^ T cells were cultured with 2.5 μg/mL anti-IL-4 (clone 11B11), 2.5 μg/mL anti-IFN-γ (clone XMG 1.2; both Bio X Cell), 10 ng/mL human IL-2, and 5 ng/mL recombinant human TGF-β1 (both PeproTech). FACS-sorted Treg cells were activated on Delta surface 96-well plates (Nunc) with 0.25 ng/mL anti-CD3 (clone 145-2C1) and 1 ng/mL anti-CD28 (clone 37.51; both Bio X Cell), 20 ng/mL recombinant human IL2 and 10 ng/mL murine IL7 for three days at a density of 2×10^6^ cells/mL.

### Seahorse extracellular flux analysis

Mitochondrial respiration and glycolytic activity of Treg cells were assessed by measuring oxygen consumption rate (OCR) and glycolytic proton efflux rate (PER), respectively, using an XFe96 extracellular flux analyzer (Seahorse Bioscience) as previously described (39). XFe96 cell culture microplates (Agilent) were pre-coated with 22 μg/mL Cell-Tak (Corning), and 1.5×10⁵ T cells per well were seeded in 1-3 technical replicates in Seahorse XF RPMI medium (Agilent) supplemented with 2 mM L-glutamine (Gibco), 1 mM sodium pyruvate and 10 mM D-glucose (both Sigma-Aldrich). Plates were incubated for 1 h at 37 °C in a CO_2_-free incubator prior to the assay. Glycolytic stress tests were performed by first measuring the basal extracellular acidification rate (ECAR), followed by addition of 0.5 μM rotenone (AdipoGen) and 0.5 μM antimycin A (Sigma-Aldrich) to inhibit mitochondrial complexes I and III, respectively. At the end of the assay, 50 mM 2-deoxy-D-glucose (2-DG) (Carl Roth) was added to completely inhibit glycolysis. To calculate the glycolytic proton efflux rate (PER) the buffer factor of the medium was included. Basal glycolysis was plotted by taking the values from the measurement right before the first injection. Compensatory glycolysis is the maximum measurement after the injection of mitochondrial inhibitors (Rotenone/Antimycin A) compared to baseline. Mitochondrial stress tests were conducted by measuring basal OCR, followed by sequential addition of 2 μM oligomycin (Biomol), 1 μM carbonyl cyanide-4-(trifluoromethoxy)-phenylhydrazone (FCCP; Biomol) and 0.5 μM rotenone plus 0.5 μM antimycin A. Basal OCR was calculated as the OCR measured before oligomycin addition minus OCR after rotenone and antimycin A treatment. Maximal OCR was calculated as the OCR following FCCP addition minus OCR after rotenone and antimycin A treatment.

### *In vitro* Treg suppression assay

CD4⁺ T cells were enriched from the spleen and LNs of female hemizygous *Slc2a3*^fl/fl^*Foxp3*^Cre/+^ mice using negative selection with the MojoSort™ Mouse CD4⁺ T Cell Isolation Kit (BioLegend). CD25^hi^ YFP⁺ and CD25^hi^ YFP^−^ T cells were subsequently FACS-sorted to obtain GLUT3-deficient and WT Treg cells, respectively. Responder T cells were isolated from spleens of congenic donor mice and labeled with CellTrace Violet (CTV; Invitrogen) according to the manufacturer’s instructions. CTV-labeled responder T cells were cultured either alone or co-incubated with WT and GLUT3-deficient Tregs at various ratios. Cells were seeded in 96-well U-bottom plates containing standard RPMI 1640 medium and stimulated with 0.5 μg/mL anti-CD3 antibody. Proliferation of CTV-labelled responder CD4^+^ and CD8^+^ T cells was assessed by flow cytometry after 4 days of culture.

### Histopathological evaluation of tissue inflammation

Tissue samples were fixed in 4 % paraformaldehyde (PFA) for 24 h, processed for paraffin embedding and sectioned at 5 μm for hematoxylin and eosin (H&E) staining, as previously described (49). H&E-stained sections were scanned using a Pannoramic 150 DX slide scanner (Sysmex). Lymphocytic infiltration and tissue inflammation in the liver, lung, colon, small intestine (SI), ear skin, pancreas, kidney and submandibular glands (SMGs) were scored as negative, mild or severe by screening the entire tissue section using the SlideViewer software (3DHistech).

### Serum cytokine quantification

To determine cytokine levels in the sera of WT and *Slc2a3*^fl/fl^*Foxp3*^Cre/Y^ mice, the LEGENDplex Mouse T Helper Cytokine Panel Version 3 (BioLegend) was used according to the following manufacturer’s instructions. Data were acquired on a BD FACSCelesta flow cytometer (BD Biosciences) with more than 3,000 bead events per sample. The resulting data were quantified against standard curves using LEGENDplex Data Analysis Software (BioLegend), and values were expressed in ng/mL.

### Quantitative real-time PCR

Total RNA was extracted using the Roti-Prep Mini Kit (Carl Roth) and cDNA was synthesized using the iScript cDNA synthesis kit (Bio-Rad). Quantitative real-time PCR was performed using the SYBR Green qPCR Master Mix (Bio-Rad) and gene-specific primers (**Table S2**). The relative transcript abundance was normalized to the expression of the housekeeping gene 18S rRNA using the 2^−ΔCT^ method as previously described (71).

### Measurement of anti-dsDNA autoantibodies

For the detection of anti-dsDNA autoantibodies, 96 well plates were coated with poly-l-lysine and incubated with 10 μg/mL calf thymus DNA (Sigma-Aldrich). After removal of unbound dsDNA, wells were blocked using 10 % FCS in PBS. Serum samples from WT and *Slc2a3*^fl/fl^*Foxp3*^Cre/Y^ mice were added in different dilutions (starting at 1:10 followed by 1:3 serial dilutions) and incubated o/n at 4 °C. After washing, 0.25 μg/mL anti-mouse IgG-Biotin or anti-mouse IgM-Biotin (both BioLegend) were added for 2 h at RT. After signal amplification using Avidin-HRP (BioLegend), 3,3’,5,5’ tetramethylbenzidine (TMB) substrate (BioLegend) was added for signal detection and absorbance at 450 nm was measured using a Flexstation III microplate reader (Molecular Devices).

### Statistical analysis

The results are presented as mean ± standard error of the means (SEM). Statistical significance between the experimental groups was assessed using an unpaired t-test, one- or two-way ANOVA, depending on the dataset using the GraphPad Prism 10 software. Sample sizes were determined based on experience and experimental complexity; however no formal tests were performed to assess the normal distribution of the data. Differences were considered significant with p < 0.05 (noted in figures as *), p < 0.01 (**) and p < 0.001 (***). Figure legends specify the number of independent samples or mice per group used in each experiment.

## AUTHOR CONTRIBUTION

Participated in research design: M.V., K.S., and M.E. Conducted experiments: M.V., M.E., X.Z. A.F., M.R. and S.M.H. Performed data analysis: M.V., S.M.H., M.E., A.F., M.R. and K.S. Wrote the manuscript: M.V. and K.S.

## ACKNOWLEDGEMENTS

We thank Dr. E. Dale Abel (Fraternal Order of Eagles Diabetes Research Center and Division of Endocrinology and Metabolism, Roy J. and Lucille A Caver College of Medicine, University of Iowa, Iowa City, IA, USA) for kindly providing *Slc2a3*^fl/fl^ mice. This work was supported by the Deutsche Forschungsgemeinschaft (DFG) SFB 1525 (“Cardio-Immune Interfaces”) – project number: 453989101; SFB-TRR 338 (“LETSimmun”) – project number: 452881907; SFB 1583 (“DECIDE”) – project number: 49262049; and individual project grants VA882/2-1 and VA882/3-2. Further support was provided by the DFG SFB 1526 (“PANTAU”), project number: 454193335; the Interdisciplinary Center for Clinical Research (IZKF) Würzburg and the Mildred Scheel Early Career Center (MSNZ) of the German Cancer Aid.

## DECLARATION OF INTERRSTS

The authors declare no competing interests.

## DATA AVAILABILITY STATEMENT

The data supporting the findings of this study are available from the corresponding author upon reasonable request.

