## Supplementary Figures S1-S6 for "The Glucose Transporter GLUT3 Controls Regulatory T Cell Function"

#### Supplementary Tables S1 and S2

### Supplementary Figure S1

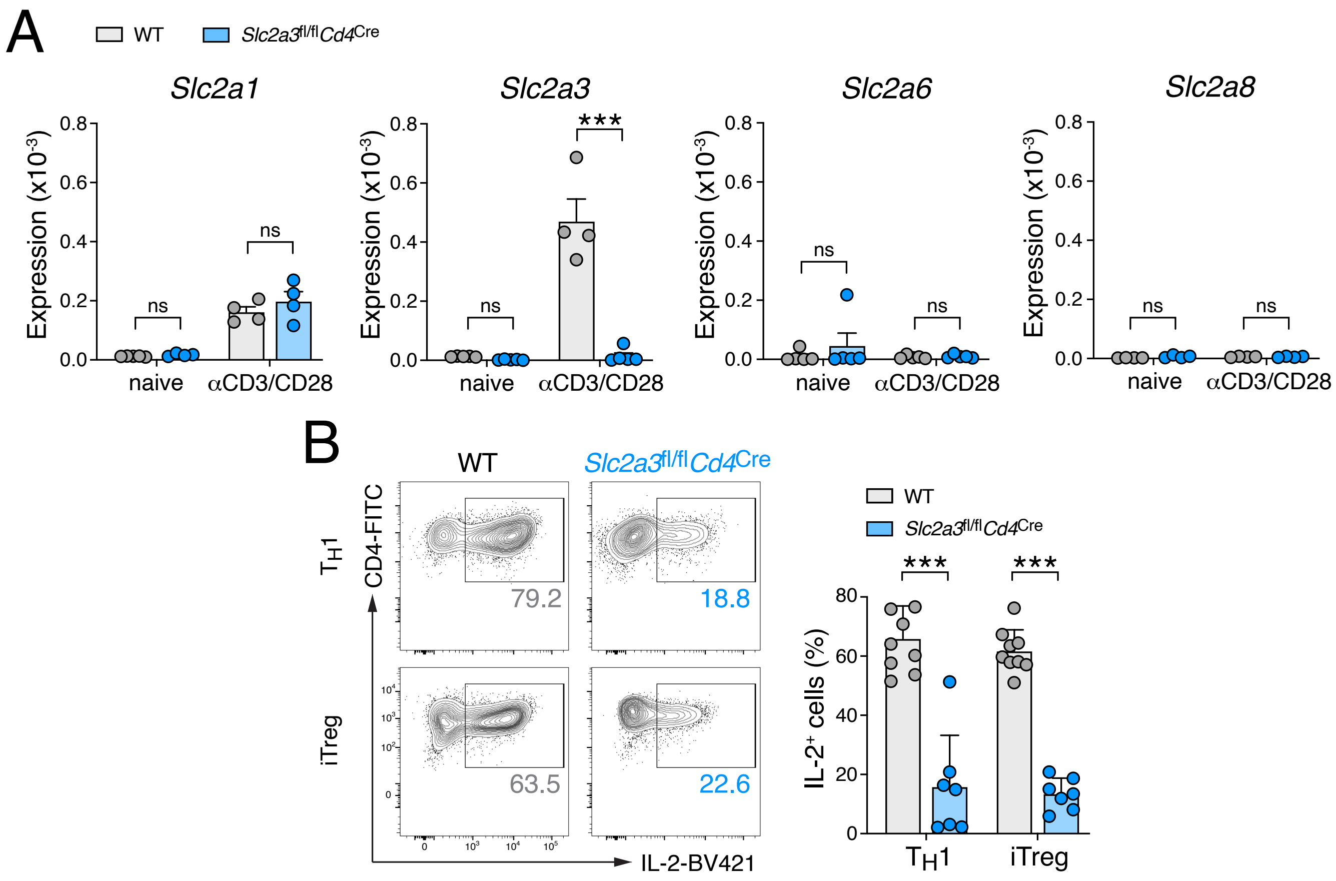


**Figure S1. Inactivation of GLUT3 in T cells impairs IL-2 production *in vitro*. (A)** Analysis of *Slc2a1* (GLUT1), *Slc2a3* (GLUT3), *Slc2a6* (GLUT6) and *Slc2a8* (GLUT8) gene expression in naïve and anti-CD3/CD28-stimulated $CD4^+$ T cells using RT-qPCR; data shown as means ± SEM of 4 mice. **(B)** Analysis of intracellular IL-2 production in *in vitro*-differentiated Th1 and iTreg cells after re-stimulation with PMA/ionomycin by flow cytometry; means ± SEM 7-8 mice. Statistical analysis in panels (A, B) was performed using two-way ANOVA. ***, $p<0.001$; ns, non-significant.

#### Supplementary Figure S2

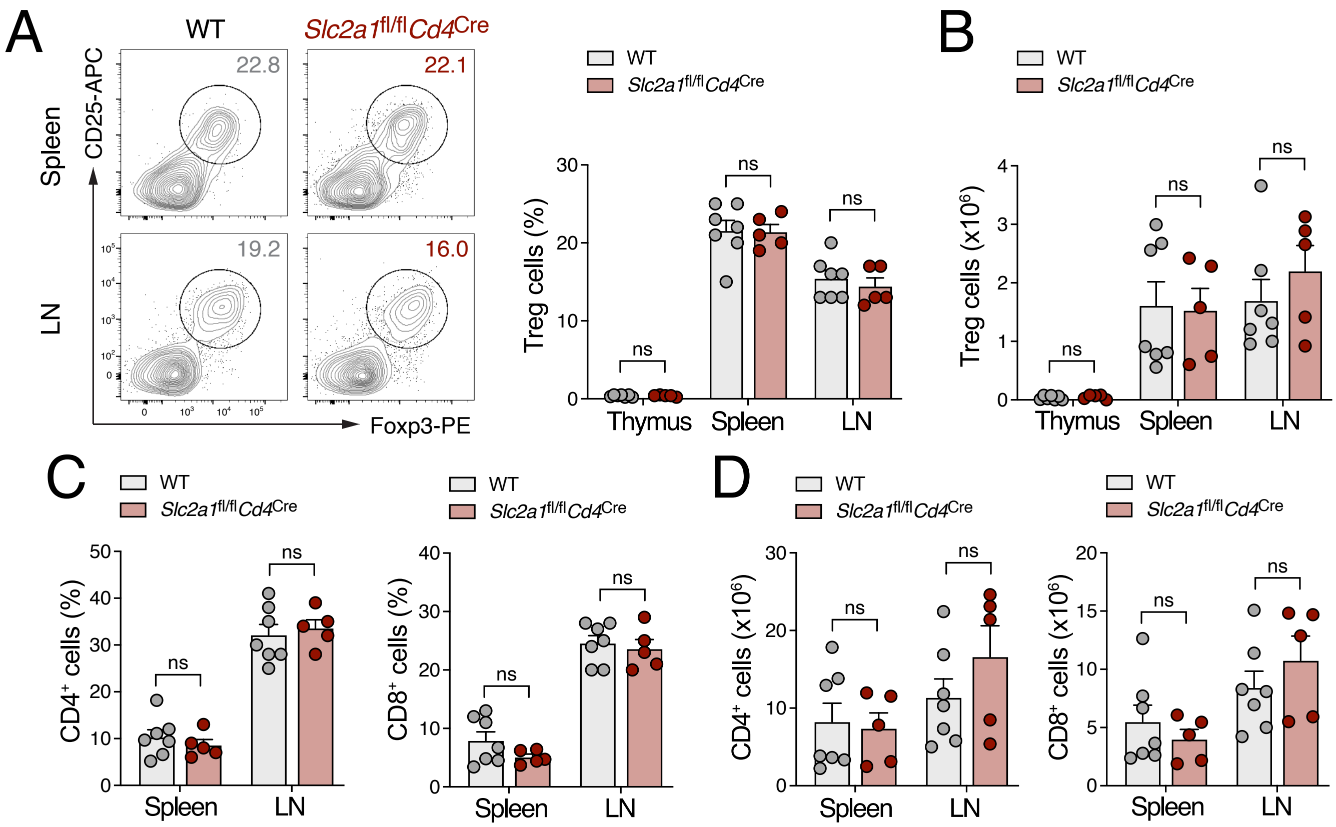


**Figure S2. Ablation of GLUT1 in T cells does not alter Treg homeostasis in lymphoid organs. (A)** Flow cytometric analysis of Foxp3$^{+}$ Treg cells in the thymus, spleen and lymph nodes (LNs) of WT and *Slc2a1*$^{fl/fl}$*Cd4*$^{Cre}$ mice; data are shown as means ± SEM of 5-7 mice. **(B)** Quantification of absolute Treg cell numbers in the indicated lymphoid organs from WT and *Slc2a1*$^{fl/fl}$*Cd4*$^{Cre}$ mice; means ± SEM of 5-7 mice. **(C and D)** Analysis of conventional CD4$^{+}$ and CD8$^{+}$ T cells in the spleen and LNs of WT and *Slc2a1*$^{fl/fl}$*Cd4*$^{Cre}$ mice. Frequencies **(C)** and absolute cell numbers **(D)** are shown as means ± SEM from 5-7 mice per group. Statistical analysis in panels (A-D) was performed using two-way ANOVA. ns, non-significant.

### Supplementary Figure S3

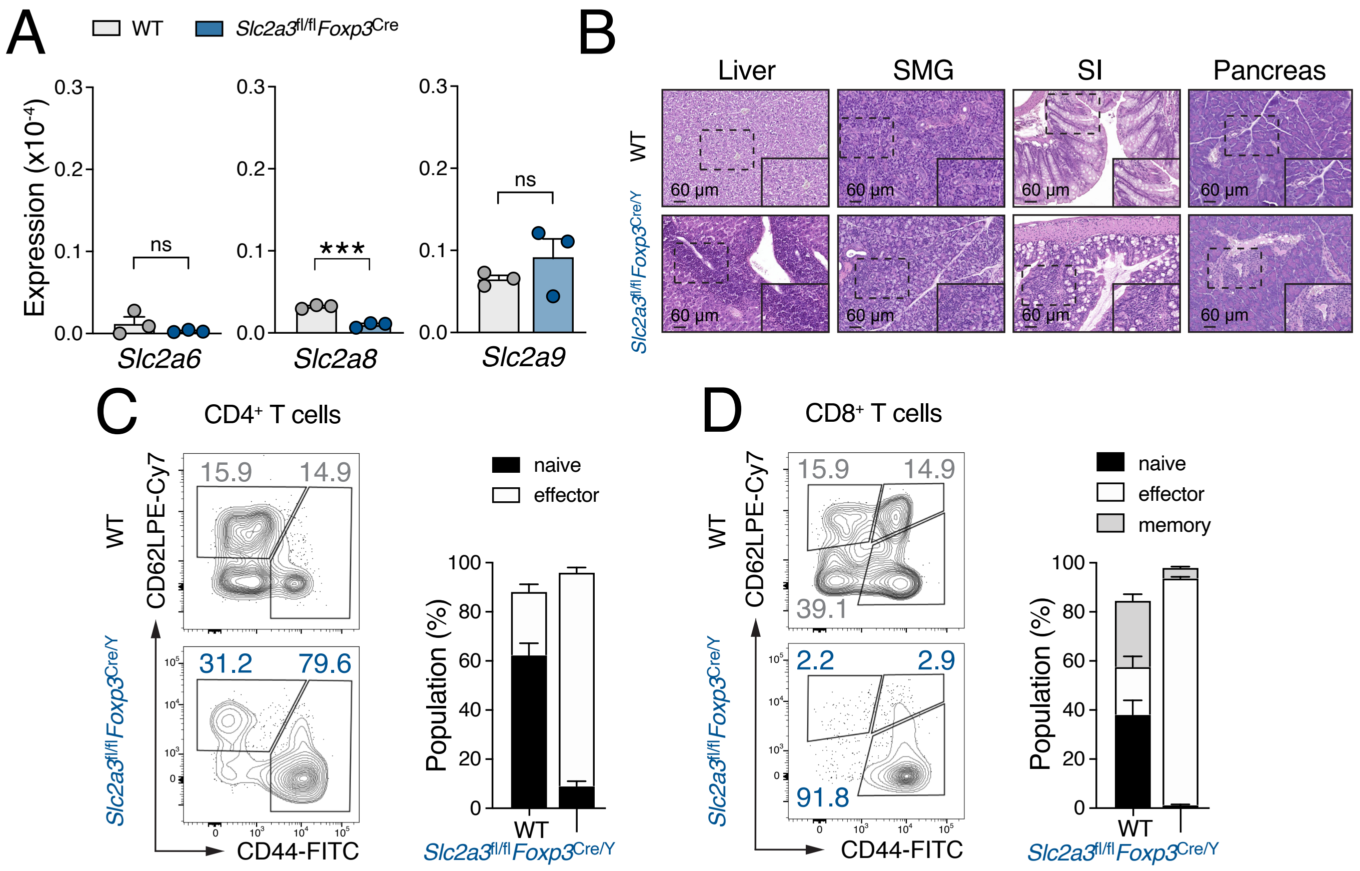


**Figure S3. Treg-specific deletion of GLUT3 causes systemic immune activation and inflammation. (A)** Analysis of *Slc2a6* (GLUT6), *Slc2a8* (GLUT8) and *Slc2a9* (GLUT9) gene expression in FACS-sorted Treg cells from WT and *Slc2a3*$^{fl/fl}$*Foxp3*$^{Cre}$ mice by RT-qPCR; data are shown as means ± SEM of 3 mice. **(B)** Representative examples of hematoxylin- and eosin-stained sections of liver, submandibular glands (SMG), small intestine (SI) and pancreas from male WT and *Slc2a3*$^{fl/fl}$*Foxp3*$^{Cre/Y}$ mice; scale bars represent 60 µm. **(C and D)** Flow cytometric analysis of naïve (CD62L$^{hi}$CD44$^{lo}$), effector (CD62L$^{lo}$CD44$^{hi}$) and central memory (CD62L$^{hi}$CD44$^{hi}$) CD4$^{+}$ **(C)** and CD8$^{+}$ **(D)** T cells in the spleen of male WT and *Slc2a3*$^{fl/fl}$*Foxp3*$^{Cre/Y}$ mice; means ± SEM of 10-13 mice. Statistical analysis in panels (A) was performed using two-way ANOVA. ***, p<0.001; ns, non-significant.

### Supplementary Figure S4

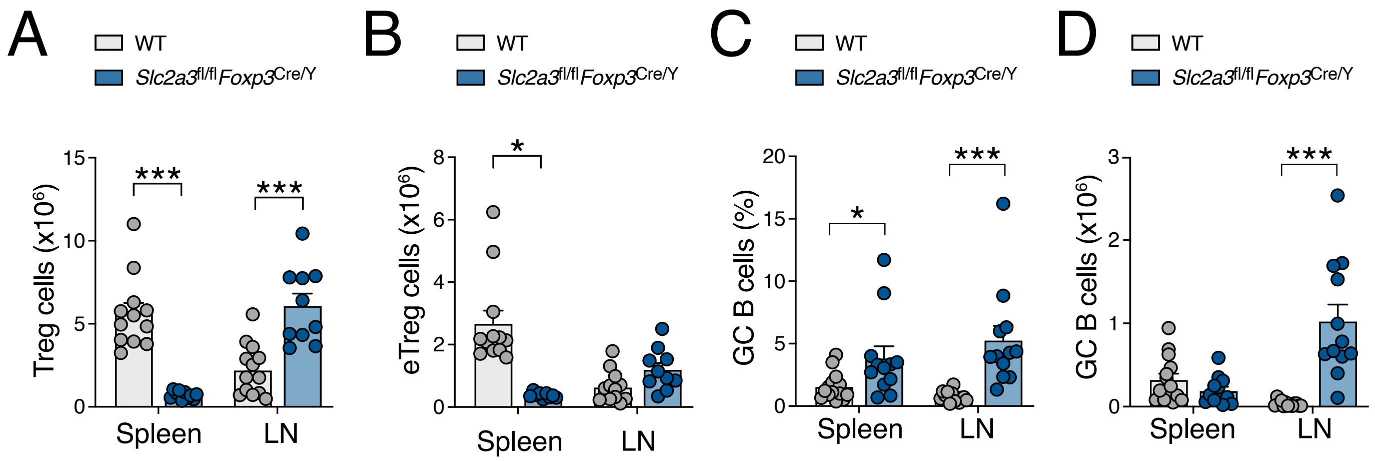


**Figure S4. GLUT3 regulates effector Treg differentiation and function. (A** and **B)** Quantification of total Foxp3$^+$ Treg cell **(A)** and effector Treg (eTreg) cell numbers **(B)** in the spleen and lymph nodes (LNs) of male WT and *Slc2a3*$^{fl/fl}$*Foxp3*$^{Cre/Y}$ mice; data are shown as means ± SEM of 10-13 mice. **(C** and **D)** Quantification of the frequencies **(C)** and total cell numbers of germinal center (GC) B cells in the spleen and LNs of male WT and *Slc2a3*$^{fl/fl}$*Foxp3*$^{Cre/Y}$ mice; data are means ± SEM of 10-13 mice. Statistical analysis in panels (A-D) was performed using two-way ANOVA. *, p<0.05; **, p<0.01, ***, p<0.001.

### Supplementary Figure S5

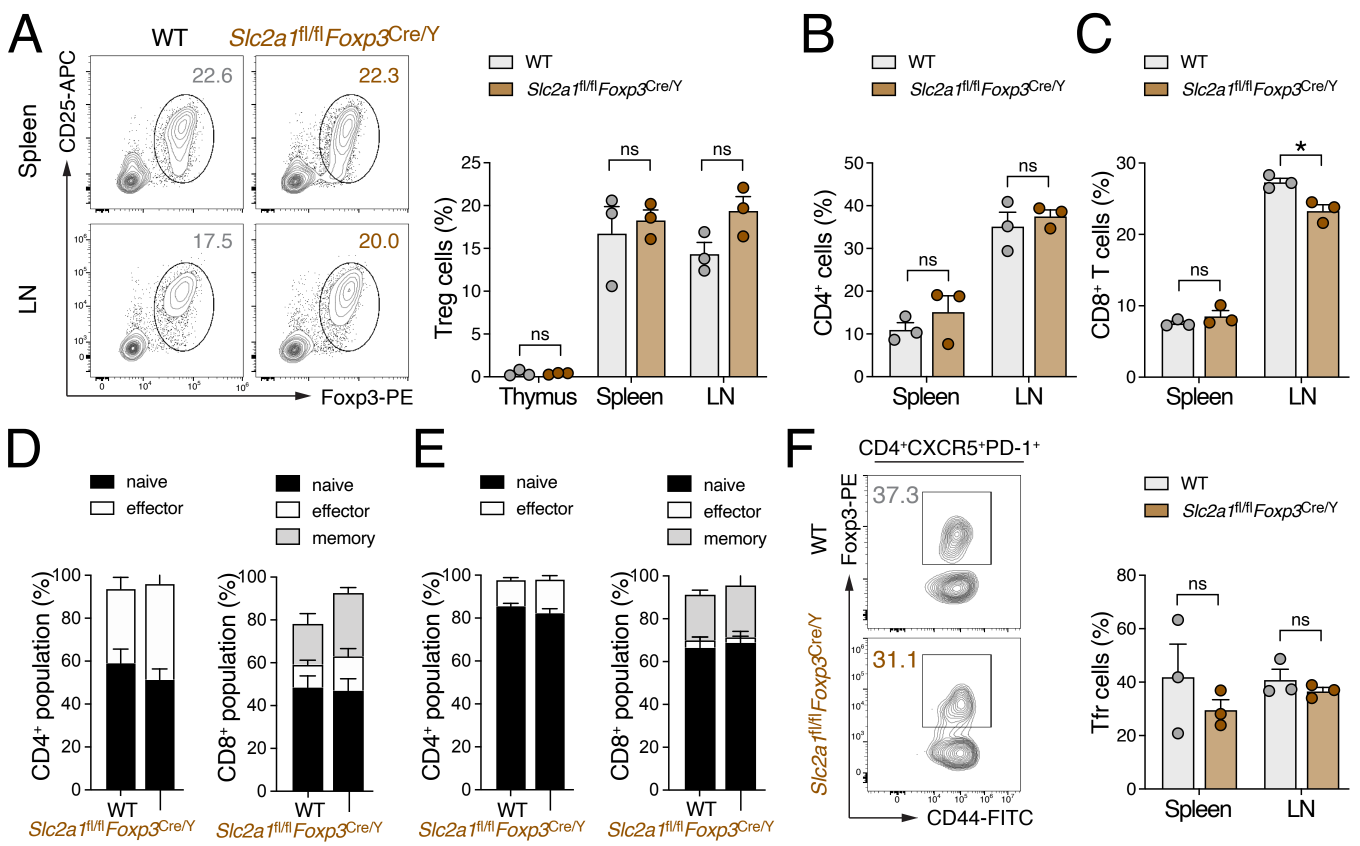


**Figure S5. Treg-specific ablation of GLUT1 does not alter Treg homeostasis, immune composition and effector differentiation. (A)** Flow cytometric analysis of Foxp3$^{+}$ Treg cells in the spleen and lymph nodes (LNs) of hemizygous male WT and *Slc2a1*$^{fl/fl}$*Foxp3*$^{Cre/Y}$ mice; data are shown as means ± SEM of 3 mice. **(B** and **C)** Analysis of conventional CD4$^{+}$ **(B)** and CD8$^{+}$ T cells frequencies **(C)** in the spleen and LNs from WT and *Slc2a1*$^{fl/fl}$*Foxp3*$^{Cre/Y}$ mice; data are shown as means ± SEM of 3 mice. **(D** and **E)** Flow cytometric analysis of naïve (CD62L$^{hi}$CD44$^{lo}$), effector (CD62L$^{lo}$CD44$^{hi}$) and central memory (CD62L$^{hi}$CD44$^{hi}$) CD4$^{+}$ and CD8$^{+}$ T cells in the spleen **(D)** and LNs **(E)** of male WT and *Slc2a1*$^{fl/fl}$*Foxp3*$^{Cre/Y}$ mice; means ± SEM of 3 mice. **(F)** Representative flow cytometric analysis of T follicular regulatory (Tfr) cells in the spleen and LNs of male WT and *Slc2a1*$^{fl/fl}$*Foxp3*$^{Cre/Y}$ mice, with quantification of Tfr cell frequencies; means ± SEM of 3 mice. Statistical analysis in panels (A-E) was performed using two-way ANOVA. *, $p<0.05$; ns, non-significant.

#### Supplementary Figure S6

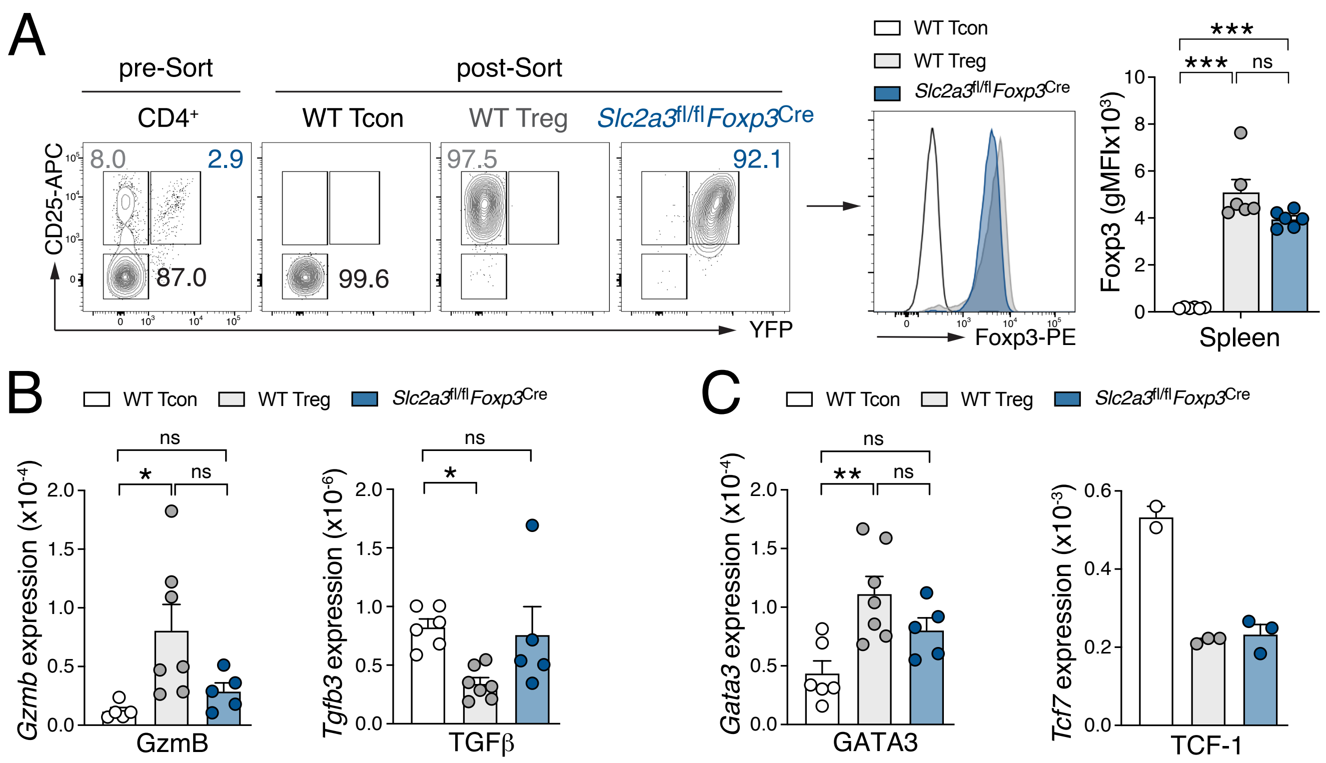


**Figure S6. Isolation and characterization of WT and GLUT3-deficient Treg cells. (A)** Representative FACS-sorting strategy to isolate T conventional (Tcon), WT and GLUT3-deficient Treg cells from female heterozygous *Slc2a3*$^{fl/fl}$*Foxp3*$^{Cre/+}$ mice (left) and verification of intracellular Foxp3 expression post sorting. **(B)** Expression of *Gzmb* (granzyme B) and *Tgfb3* (TGFβ) in FACS-sorted Tcon, WT and GLUT3-deficient Treg cells from female heterozygous *Slc2a3*$^{fl/fl}$*Foxp3*$^{Cre/+}$ mice measured by RT-qPCR; data shown as means ± SEM of 5-7 mice. **(C)** Expression of *Gata3* (GATA-3) and *Tcf7* (TCF-1) in FACS-sorted Tcon, WT and GLUT3-deficient Treg cells from female heterozygous *Slc2a3*$^{fl/fl}$*Foxp3*$^{Cre/+}$ mice measured by RT-qPCR; data shown as means ± SEM of 2-7 mice. Statistical analysis in panels (A) was performed using two-way ANOVA. *, p<0.05; **, p<0.01; ***, p<0.001; ns, non-significant.

**Table S1. List of antibodies used in this study**

| Antibody | Clone | Company | Cat# | RRID | Dilution |
|---|---|---|---|---|---|
| anti-B220 (FITC conjugated) | RA-6B2 | BioLegend | 103206 | AB_312991 | 1:400 |
| anti-CD3 (PE conjugated) | 145-2C11 | BioLegend | 100308 | AB_312673 | 1:400 |
| anti-CD3 (FITC conjugated) | 145-2C11 | BioLegend | 100306 | AB_312671 | 1:400 |
| anti-CD4 (BV421 conjugated) | RM4-5 | BioLegend | 100544 | AB_2563052 | 1:400 |
| anti-CD4 (BV605 conjugated) | GK1.5 | BioLegend | 100451 | AB_2564591 | 1:400 |
| anti-CD8 (BV711 conjugated) | 53-6.7 | BioLegend | 100748 | AB_2563510 | 1:400 |
| anti-CD11b (PE conjugated) | M1/70 | BioLegend | 101208 | AB_312791 | 1:400 |
| anti-CD19 (BV421 conjugated) | 6D5 | BioLegend | 115538 | AB_11203527 | 1:500 |
| anti-CD25 (APC conjugated) | PC62 | BioLegend | 102012 | AB_312861 | 1:400 |
| anti-CD38 (APC conjugated) | 90 | BioLegend | 102712 | AB_312932 | 1:400 |
| anti-CD44 (FITC conjugated) | IM7 | BioLegend | 103043 | AB_2561391 | 1:400 |
| anti-CD45 (BV510 conjugated) | 104 | BioLegend | 109838 | AB_2561393 | 1:400 |
| anti-CD62L (PE-Cy7 conjugated) | MEL-14 | BioLegend | 104418 | AB_313102 | 1:400 |
| anti-CXCR5 (PerCP-Cy5.5) | SPRCL5 | eBioscience | 46-7185-82 | AB_2561972 | 1:100 |
| anti-Foxp3 (PE conjugated) | FJK-16s | ThermoFisher Scientific | 12-5773-82 | AB_465936 | 1:200 |
| anti-Foxp3 (FITC/AF488 conjugated) | FJK-16s | ThermoFisher Scientific | 53-5773-82 | AB_763537 | 1:200 |
| anti-Foxp3 (APC conjugated) | FJK-16s | ThermoFisher Scientific | 17-5773-82 | AB_469457 | 1:200 |
| anti-F4/80 (BV421 conjugated) | BM8 | BioLegend | 123137 | AB_2563102 | 1:400 |
| anti-GFP (AF488 conjugated) | n/a | ThermoFisher Scientific | A-21311 | AB_221477 | 1:100 |
| anti-GM-CSF (PE conjugated) | MP1-22E9 | BioLegend | 505406 | AB_315382 | 1:200 |
| anti-GL7 (FITC/AF488 conjugated) | GL7 | BioLegend | 144612 | AB_2563285 | 1:200 |
| anti-IL-2 (BV421 conjugated) | JES6-5H4 | BioLegend | 503826 | AB_2650897 | 1:200 |
| anti-NK1.1 (FITC/AF488 conjugated) | PK136 | BioLegend | 108717 | AB_493183 | 1:200 |
| anti-PD1 (APC conjugated) | RMP1-39 | BioLegend | 109112 | AB_10613470 | 1:400 |
| anti-Siglec F (APC/AF647 conjugated) | E50-2440 | BD Bioscience | 562680 | AB_2687570 | 1:400 |

**Table S2. List of RT-qPCR primers used in this study.**

| Target | Forward | Reverse |
|---|---|---|
| *18S* | CGGCGACGACCCATTCGAAC | GAATCGAACCCTGATTCCCCGT |
| *Slc2a1* | GAGACCAAAGCGTGGTGAGT | GCAGTTCGGCTATAACACTGG |
| *Slc2a3* | ATCGTGGCATAGATCGGTTC | TCTCAGCAGCTCTCTGGGAT |
| *Slc2a6* | GCCCAGGAGGTCATTGAGTA | CCAGCACTACACCTGGACAA |
| *Slc2a8* | GGCATGTAGCACATGAGCAG | GGCATCCTCCTGGCCTAT |
| *Slc2a9* | TTGCTTTAGCTTCCCTGATGTG | GAGAGGTTGTACCCGTAGAGG |
| *Gzmb* | CCACTCTCGACCCTACATGG | GGCCCCCAAAGTGACATTTATT |
| *Tgfb3* | TCTCCACTGAGGACACATTGA | ATTCGACATGATCCAGGGAC |
| *Gata3* | AGGATGTCCCTGCTCTCCTT | GCCTGCGGACTCTACCATAA |
| *Tcf7* | AGCTTTCTCCACTCTACGAACA | AATCCAGAGAGATCGGGGGTC |
| *Il10* | TGTCAAATTCATTCATGGCCT | ATCGATTTCTCCCCTGTGAA |
| *Ebi3* | GGGATGCCAGAGCACCTG | TCCTAGCCTTTGTGGCTGAG |
| *Ctla4* | TTTTGTAGCCCTGCTCACTCT | CTGAAGGTTGGGTCACCTGTA |
| *Hif1a* | AAACTTCAGACTCTTTGCTTCG | CGGCGAGAACGAGAAGAA |
| *Irf4* | CAAAGCACAGAGTCACCTGG | TGCAAGCTCTTTGACACACA |
| *Prdm1* | TTCTCTTGGAAAAACGTGTGGG | GGAGCCGGAGCTAGACTTG |
